# A 3D Printed Ventilated Perfused Lung Model Platform to Dissect the Lung’s Response to Viral Infection in the Presence of Respiration

**DOI:** 10.1101/2023.09.24.559194

**Authors:** I. Deniz Derman, Mecit Altan Alioglu, Dishary Banerjee, Sarah E. Holton, Danielle Nicole Klunk, Momoka Nagamine, Syed Hasan Askari Rizvi, Carmen Mikacenic, Nazmiye Celik, Diana Cadena Castaneda, Warang Prajakta, Phylip Chen, Michael Schotsaert, Mark E. Peeples, Karolina Palucka, Jonathan Koff, Ibrahim T. Ozbolat

## Abstract

In this study, we developed a three-dimensionally (3D) printed lung model that faithfully recapitulates the intricate lung environment. This 3D model incorporated alveolar and vascular components that allow for a comprehensive exploration of lung physiology and responses to infection *in vitro*. In particular, we investigated the intricate role of ventilation on formation of the alveolar epithelial layer and its response to viral infections. In this regard, we subjected our 3D printed, perfused lung model to a continuous respiratory cycle at the air-liquid interface (ALI) for up to 10 days followed by infection with two viruses: influenza virus (Pr8) and respiratory syncytial virus (RSV), at two different concentrations for 24 or 48 h. The results revealed that ventilation induced increased tight-junction formation with better epithelial barrier function over time, facilitated higher expression of alveolar epithelial specific genes, enabled higher level of infection with an increased progression of viral spread and replication over time, and modulated the production of pro-inflammatory cytokines and chemokines. Our findings represent a critical step forward in advancing our understanding of lung-specific viral responses and respiratory infections in response to ventilation, which sheds light on vital aspects of pulmonary physiology and pathobiology.

## INTRODUCTION

Studying lung-specific responses to infection has been challenging due to the limited availability of human lung tissue(s) and limitations of animal models, which often fail to fully recapitulate human physiology (*1*, *2*). To bridge this gap and to gain a better understanding of viral infections, the development of robust in-vitro human lung models is crucial. Over the years, researchers have made significant progress in developing increasingly simple to complex lung tissue models that reproduce specific components of human lung responses (*3–6*). However, all these models have limitations, particularly to capture complex cell-cell and cell-matrix interactions and to replicate the intricate 3D anatomy of the lung parenchyma (*7*, *8*). Creating a sophisticated 3D in-vitro lung model presents several challenges due to the complex anatomy of the lung matrix and parenchyma, which include the requirement for connections between alveolar units and airways surrounded by endothelium (*9–12*). Although lung tissue slices offer an important approach for identifying immune and physiological events in the lung, they are limited by the difficulty to obtain human samples, limitations on longitudinal experiments, and challenges in modeling host-pathogen interactions (*13*, *14*). In the past decade, lung-on-a-chip platforms have been developed to recapitulate the behavior of the human lung under mechanical strain and perfusion. While these platforms offer significant advancements, they often lack the incorporation of relevant lung compartments within an anatomical configuration, like the human lung and surrounded by a vascular compartment also limited by matrix choices, which change effects on lung epithelium (*15–18*). For instance, in-vitro models of lung have focused on reconstructing alveolar-like structures using primary alveolar type II cells in 3D collagen gels, which supports the production of surfactant proteins (*19*, *20*). Similarly, in-vitro tissue models of human bronchial epithelium have been useful for studying cellular interactions in chronic obstructive lung diseases (*21–23*). Co-culture systems of distal lung epithelial cells and microvascular endothelial cells have also allowed investigation of cellular interactions at the alveolar-capillary barrier during acute lung injury (*24*, *25*). However, these models do not capture the complex 3D anatomy of lung parenchyma and pulmonary circulation (*26–32*).

The core function of the lung, to exchange oxygen and carbon dioxide, hinges on the harmonious interaction of ventilation and perfusion(*33*). Recent advancements in technologies such as 3D printing have ignited a new era in respiratory research because 3D printing offers an array of advantages, from the precise fabrication of complex structures with accurate cell patterning to the localized deposition of cells, which effectively mimic the intricate lung microenvironment. This technique not only streamlines manufacturing but also enables the creation of multiple platforms in parallel, which contribute to medium/high-throughput solutions for respiratory research (*34*). This synergy of cutting-edge methodologies sets the stage for an advanced understanding of lung-specific immune responses for respiratory diseases. Through the faithful replication of the intricate 3D anatomy and dynamic cellular interactions, these models provide a physiologically-relevant environment for the exploration of infectious diseases. Furthermore, this approach may empower studies into the dynamics of lung-immune interactions, which may aid the development of novel therapeutic strategies.

To address the constraints posed by current models, we proposed harnessing the potential of 3D printing to craft a ventilated, perfused lung model. The dynamic growth capacity of our 3D lung model permits continuous observation of cellular behavior, which offers insights previously unattainable by conventional *in-vitro* systems. By combining these technological advancements with studies of viral infection, a comprehensive understanding of lung-specific immune responses and the complexity of respiratory infections is within reach. Here, we focused on the role of ventilation-induced mechanical forces on the formation of the alveolar epithelial layer and its response to the viral infection in the 3D printed lung model. To comprehensively study this, these experiments utilized two different viruses: the influenza virus (Pr8) and respiratory syncytial virus (RSV). By varying experimental conditions, which included different viruses, infection duration, and viral concentrations, we aimed to gain a thorough understanding of how this 3D lung model responds to these pathogens. Over 10 days with continuous respiration at the air-liquid interface (ALI), we found that alveolar epithelial cells had improved tight-junction formation, epithelial barrier function and expressed more of alveolar epithelial-specific genes. More importantly, ventilated samples exhibited higher levels of viral infection with modulated production of pro-inflammatory cytokines and chemokines. Overall, our 3D printed lung model, designed to mimic the intricate lung environment, has yielded novel insights into the lung response to viral infections in the presence of respiration.

## RESULTS

### Characterization of the Ink

Before investigating the 3D printing of the lung model, we investigated the rheological properties of the fugitive inks, where all Pluronic concentrations showed shear thinning behavior since the viscosity of samples decreased as the shear rate increased (**Figure S1A**). As the concentration of Pluronic increased, the viscosity of the inks increased. The addition of 1.5% PEO did not alter the viscosity significantly. To determine the frequency dependence of the ink, frequency sweep tests were carried out within the linear viscoelastic region. All samples exhibited G’ > G’’ over the entire range (**Figure S1B**). The amplitude sweep test, as depicted in **Figures S1C-D**, showed the viscoelastic behavior of inks. The results demonstrated a linear viscoelastic region in the gel state, where G’ > G’’. Also, the inclusion of PEO reduced the storage modulus of the Pluronic ink, as has been reported previously (*41*). Since fugitive inks were subjected to high shear stress during printing (but not after printing), self-recovery properties were investigated by a thixotropic recovery test (**Figure S1E**). All samples swiftly regained their viscosity upon testing, reverting to their original consistency within a matter of seconds. The fugitive inks, as anticipated, exhibited the expected properties of yield stress gels, demonstrating their ability to maintain their structural integrity under stress. Pluronic has been widely utilized as a fugitive ink due to its ease of use and ability to liquefy at 4 °C (*42*). Initially, 34, 38, and 42% (w/v) Pluronic F-127 were tested as a fugitive ink. With higher concentrations, better attachment to the gelatin surface was achieved and printed filaments were less likely to move during printing. With 38% (w/v) concentration, we were able to obtain stable printing performance without filament displacement. Therefore, 38% Pluronic was utilized as the fugitive ink for the remaining experiments. Moreover, we optimized our fugitive ink further by adding high-molecular-weight PEO to mitigate the viscous-fingering phenomenon (*43*). The printability of fugitive inks was also investigated using a testbed, which replicated the conditions experienced during lung model printing, where fugitive inks were printed on a crosslinked 10% gelatin surface **(Figure S2A)**. With 38% Pluronic F-127, we were able to achieve stable performance and the addition of PEO did not change printability (**Figures S2B**).

### Lung Model Fabrication

Upon optimizing ink properties and the 3D printing process, lung model constructs were fabricated with precise reconstitution of the alveolar compartment, airway, and perfusion channel for pulmonary circulation **(Figures 1 and S3)**. The fabricated lung model displayed anatomical and physiological features, which faithfully reproduce the 3D anatomy and vascular elements of the lung, although in a simplified form. The lung model was effectively connected to the ventilation and perfusion systems, enabling a continuous respiratory cycle (3-sec inhalation and 3-sec exhalation) and cell medium perfusion. The ventilation system provided humidified air with controlled pressure (± 1-1.5 cm H_2_O) to mimic physiological breathing conditions (*44*, *45*). The perfusion system maintained at a constant flow rate to ensure nutrient supply and waste removal for the cultured cells **(Figures 1B-C and Video S1).**

**Figure 1:**
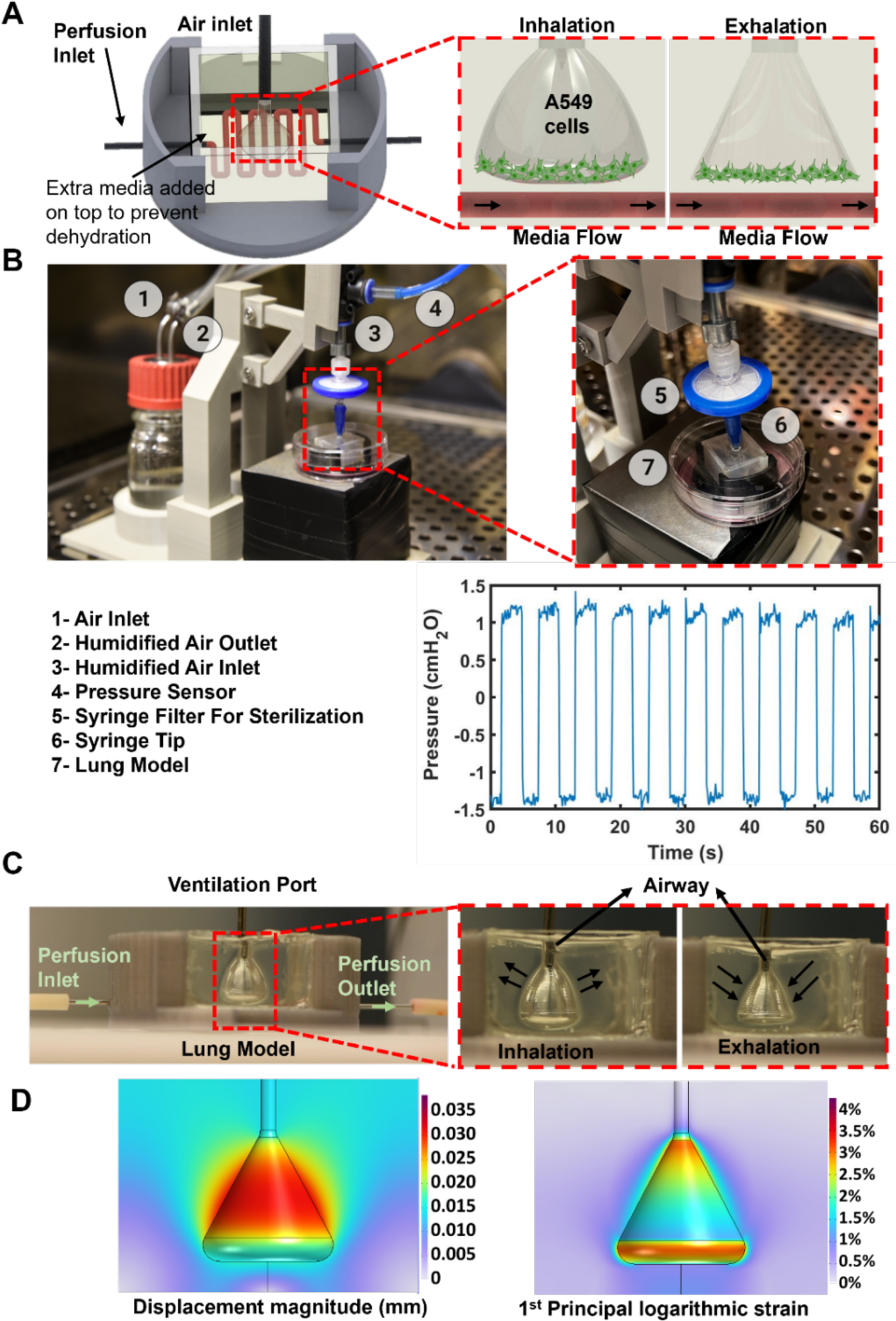
**(A)** The schematic diagram of the lung model. **(B)** The device set up inside an incubator, where air supplied from an air pump was humidified in a humidification chamber, sterilized with a syringe filter, and directed into the lung model. The air pump pressure was adjusted to achieve a ±1.5 cm H_2_O breathing range, **(C)** enabling inhalation and exhalation of the lung model. **(D)** Elastic deformation of the ventilated lung was modeled to demonstrate displacement magnitude and strain.

Experiments were conducted to gain insights into the mechanical behavior of the 3D lung model under dynamic breathing conditions, particularly in terms of displacement and strain distribution. The displacement field analysis revealed that the sides of the alveolar lung cavity experienced greater displacement compared to the bottom **(Figure 1D)**. However, the strain plot indicated that the bottom of the lung experienced more strain than the rest of the structure. This discrepancy suggested that while the sides may have shifted more, the overall displacement of the bottom was constrained, possibly due to rigid body motion or structural stability mechanisms (*46*, *47*). As a result of these simulation findings, a practical adjustment was made in the experimental setup. A549 cells were specifically seeded at the bottom of the alveolar chamber, where the displacement and strain were relatively less prominent. This strategic placement aimed to ensure that the cellular environment remained relatively stable, minimizing a possible substantial impact of displacement during the experimental procedures. Moreover, to maintain the viability of cells and facilitate ALI, perfusion channels were exclusively positioned at the bottom of the chamber. This arrangement aimed to provide a consistent supply of nutrients and facilitate the removal of waste products, enhancing the overall cellular health and enabling the accurate study of lung dynamics without compromising cellular viability. In essence, the simulation experiments provided crucial insights into the mechanical dynamics of the 3D lung model during breathing-like motions. The information obtained from these simulations guided the experimental design and placement of cells to optimize the accuracy and reliability of the study’s outcomes.

Next, A549 human adenocarcinoma cells were successfully cultured in the lung model under ALI conditions. We classified our experimental conditions into four distinct groups: Ventilation (-)/Perfusion (-), Ventilation (+)/Perfusion (-), Ventilation (-)/Perfusion (+) and Ventilation (+)/Perfusion (+). These groups were devised to encompass a spectrum of physiological scenarios and allow us to compare the individual and combined impacts of ventilation and perfusion on our lung model’s performance.

Immunofluorescent staining of the cultured cells revealed the expression of ZO-1 (**Figures 2A and S4**), a tight-junction protein, which indicates the formation of tight junctions and barrier integrity at Days 4, 7 and 10. To evaluate the integrity of tight junctions, we quantitatively assessed the distance between tight junctions at Day 10 (**Figure 2B)**. Immunofluorescent images revealed an increase in tight junction density over time. While we conducted Zonula Occludens-1 (ZO-1) staining across all experimental groups, it is important to note that ZO-1-to-ZO-1 distance measurements were not performed for the Ventilation (-)/Perfusion (-) and Ventilation (+)/Perfusion (-) groups as these groups did not develop well-defined (measurable) tight junctions. The Ventilation (+)/Perfusion (+) group exhibited increased tight junction formation with a smaller ZO-1-to-ZO-1 distance compared to the Ventilation (-)/Perfusion (+) group. To further confirm the barrier function (the integrity and tightness of the epithelial barrier), transepithelial electrical resistance (TEER) measurements over the 10 days of ALI culture were completed (**Figure 2C**). After 10 days of ALI culture, the mean TEER value for the Ventilation (-)/Perfusion (-) group measured ∼ 16 Ω × cm^2^, which was significantly lower than the average TEER value of ∼ 43 Ω × cm^2^ observed in the Ventilation (+)/Perfusion (+) group (p = 0.0003). Notably, all groups displayed an increasing trend in TEER values over time. To further confirm the tightness of the epithelial barrier, we conducted a permeability study using 4 kDa fluorescein isothiocyanate (FITC)-labeled Dextran (**Figures 2D and S5**). The results revealed a significant reduction in the diffusion of Dextran from the Ventilation (-)/Perfusion (-) group to the Ventilation (+)/Perfusion (+) group, where the acellular alveolar chamber was used as a control. These results demonstrated the dynamic nature of the epithelial tissue formation with a more functional barrier formation over time. As the functional analysis revealed measurable tight junctions for Perfusion (+) groups, which could be due to the fact that Perfusion (-) groups did not facilitate an appropriate environment for A549 cell growth, we proceeded with the further investigation of Ventilation (+)/Perfusion (+) and Ventilation (-)/Perfusion (+) groups for the rest of this study.

**Figure 2:**
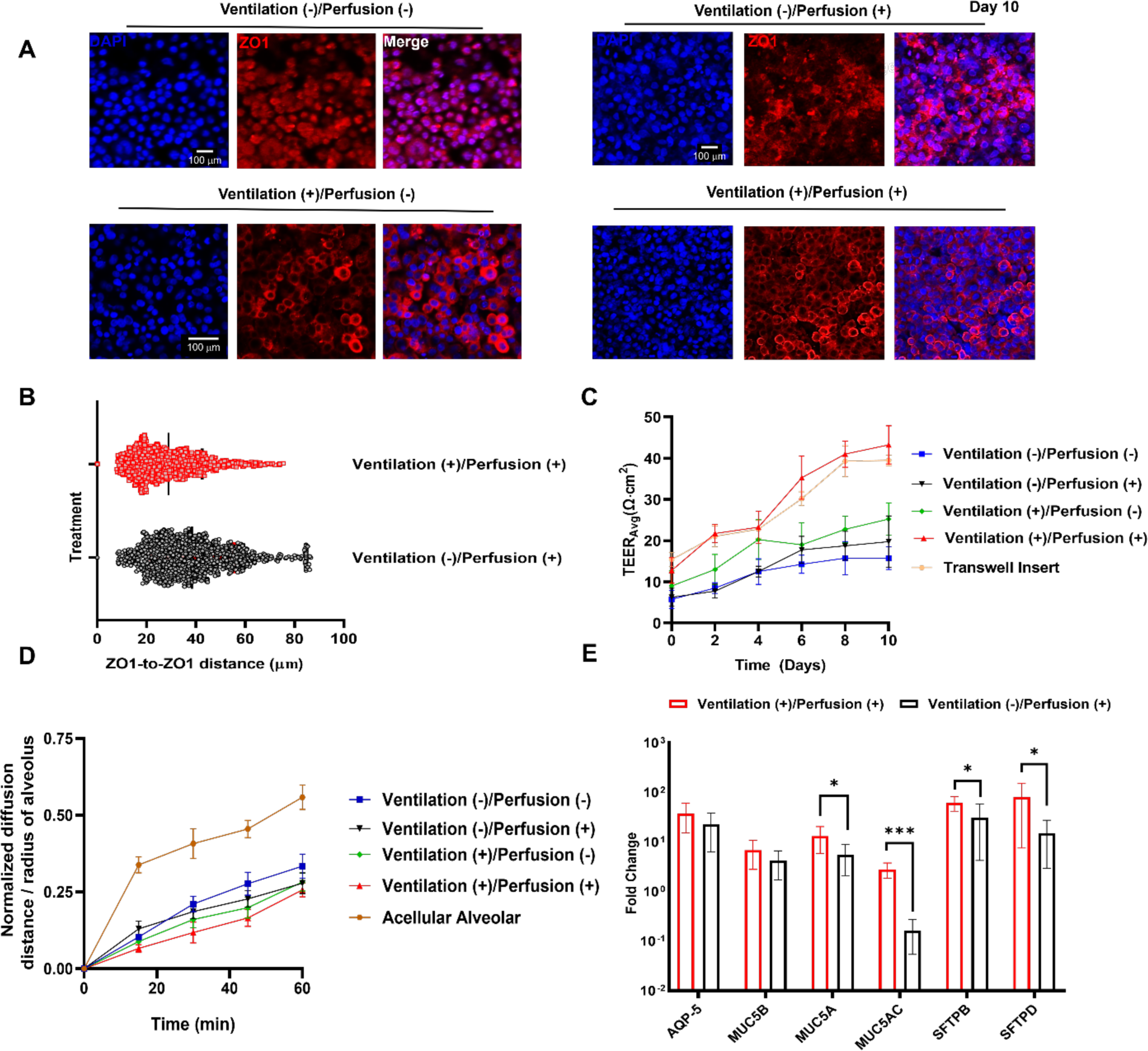
**(A)** Evaluation of tight junctions of A549 cells in four different groups using ZO-1 staining at Day 10. **(B)** Characterization of ZO-1 staining with respect to ZO-1-to-ZO-1 distance on Day 10. **(C)** Measurement of TEER values during the 10-day ALI culture. **(D)** Comparison of Dextran permeability and **(E)** evaluation of gene expressions via qRT-PCR (*n=*3; *p*<*0.05 and *p***<*0.001).

Within the context of our investigation, the lung model also supported the differentiation of A549 cells. This was substantiated by the discernible presence of MUC5B expression (**Figure S6**) and the discernment of cilia through α-tubulin expression (**Figure S7**). Based on fluorescent images, we observed no significant difference in MUC5B expression between Ventilation (+)/Perfusion (+) and Ventilation (-)/Perfusion (+) groups. However, when assessing α-tubulin expression, a notable discrepancy emerged between the two groups, with stronger expression of α-tubulin observed in the Ventilation (+)/Perfusion (+) group. These findings highlight the role of ventilation on fostering respiratory epithelial cell differentiation and function in the 3D printed lung model. These results demonstrated that the 3D lung model provided a conducive microenvironment for the differentiation and function of respiratory epithelial cells. Building upon this premise, we further delved into the assessment of expression levels pertaining to key genes, including aquaporin-5 (AQP-5), mucin-5B (MUC5B), mucin-5A (MUC5A), mucin-5AC (MUC5AC), surfactant protein-B (SFTPB), and surfactant protein-D (SFTPD) **(Figure 2E)**. Although there was no significant difference between non-ventilated and ventilated groups in AQP-5 and MUC5B expression at Day 10, there was a significant difference between Ventilation (-)/Perfusion (+) and Ventilation (+)/Perfusion (+) groups in the expression levels of MUC5A (∼2-fold increase), MUC5AC (∼17-fold increase), SFTPB (∼2-fold increase), and SFTPD (∼5-fold increase) at Day 10.

### Viral Infection

After confirming the function of the lung model, we sought to determine whether this model allowed for the study of influenza virus and/or RSV. Influenza virus (A/Pr8/34) expressing mCherry or RSV-A2 expressing RFP at an inoculating dose of 1×10^5^ or 1×10^6^ PFU, respectively, were exposed at the apical surface of the ALI cultures in serum-free Pneumacult ALI medium. As a mock control, ALI cultures were also exposed to the medium without virus. At 24 or 48 h post inoculation, ALI cultures were collected and Calcein (violet) staining was performed. RFP signal indicated active infection, as RFP served as a gene reporter for the nonstructural protein 1 (NS1) for each virus. At 24 h post Pr8 infection, the Ventilation (+)/Perfusion (+) group showed a significantly increased average proportion of RFP^+^ infected cells at 1×10^5^ PFU (57%) compared to samples from the Ventilation (-)/Perfusion (+) group (∼23% for 1×10^5^ PFU, ∼ 34% for 1×10^6^ PFU) group (**Figure 3**). Interestingly, at 48h of Pr8 infection, the Ventilation (+)/Perfusion (+) group had a significantly increased proportion of RFP^+^ cells at 1×10^5^ PFU (∼53%) and 1×10^6^ PFU (40%) compared to the samples from the Ventilation (-)/Perfusion (+) group (∼25% for 1×10^5^ PFU, ∼39% for 1×10^6^ PFU). Based on the Pr8 infection, we observed a significant difference between ventilated and non-ventilated samples, particularly evident within the first 24 h of Pr8 infection. However, at the end of the 48-h infection period, there was no significant difference between Ventilation (+)/ Perfusion (+) and Ventilation (-)/ Perfusion (+) groups (**Figure 3**). Intriguingly, our investigation into cell viability using Calcein staining (**Figures 3A2-B2 and S8-S12**) showed a significant decrease in cell viability observed at the conclusion of the 48 h infection period. This decrease in cell viability was more pronounced in the higher-level viral infections. In contrast, lower infection rates, such as those in the RSV infections, may mitigate the impact of ventilation-induced mechanical forces, resulting in comparatively higher cell viability.

**Figure 3:**
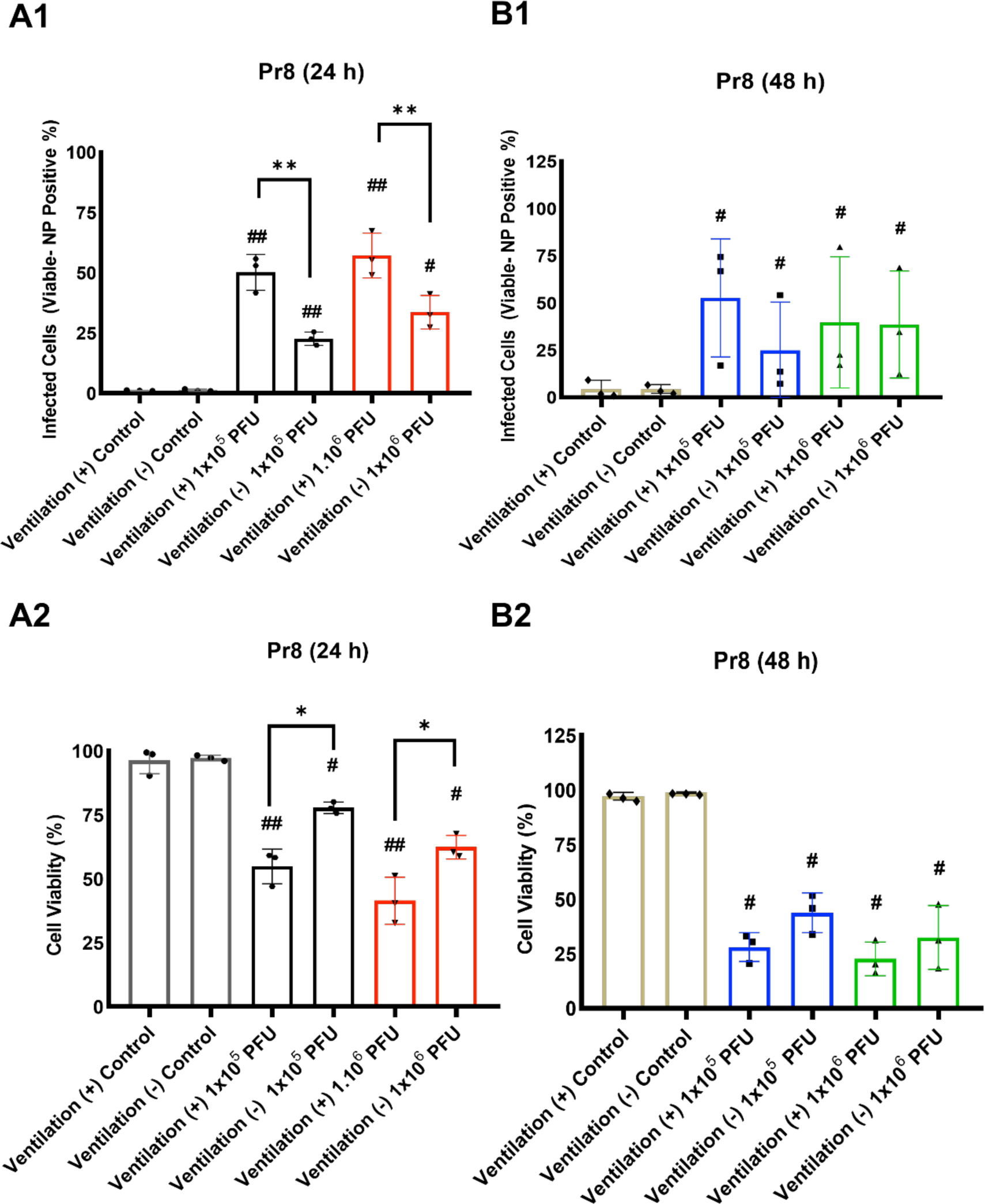
Quantification of Pr8 infection based on the Calcein and nucleoprotein (NP) positive signal. **(A1 and B1)** Flow cytometry based on infected cells (%) at 24 or 48 h and, **(A2 and B2)** cell viability (%) after 24 or 48 h of the viral infection (*n=*3; *p*<*0.05, *p**<*0.01 and *p***<*0.001; *p*^#^ <0.05 and *p*^##^ <0.01 indicate significance with respect to the control of the same group).

For the RSV infection, at the end of 24 h, the Ventilation (+)/Perfusion (+) groups showed an increased presence of RFP signal (1.8% for 1×10^5^ PFU, 4.6% for 1×10^6^ PFU) compared to the samples from the Ventilation (-)/Perfusion (+) group (1.6% for 1×10^5^ PFU, 1.7% for 1×10^6^) although these results were not being statistically significant (**Figure 4**). Interestingly, our study revealed a substantial discrepancy in infection rates between the RSV and Pr8, where RSV showed a considerably lower infection rate. This highlights the known slower viral kinetics of RSV infection in epithelial cells compared to influenza (*48*). This contrast in infection dynamics between the two viruses underscores the virus-specific variability in infectivity and highlights the importance of considering the distinct behavior of respiratory viruses when studying their interactions with lung models. Furthermore, when comparing the proportion of RSV-infected cells at 24 h and 48 h, we found that the infection rates had generally increased by 48 h, which indicates the progression of viral spread and replication over time. In the Ventilation (+)/Perfusion (+) group, while infection rates were higher at 48 h, it is important to note that these differences were not statistically significant compared to the 24 h infection rates. This observation suggests that the infection dynamics in the Ventilation (+)/Perfusion (+) groups plateaued between the 24 h and 48 h time points, which potentially reached a saturation of infection. The observed differences in infection dynamics between the Pr8 and RSV viruses also align with the viability data obtained in our study. Pr8 infection caused a decrease in the number of infected cells between 24 h and 48 h. This decrease in infected cell numbers suggests a potential phenomenon of cell death or viral-induced cytopathic effects over time. These insights were corroborated with the viability data, which indicated a significant decrease in cell viability after 48 h of infection. Therefore, the observed decrease in the number of infected cells in the Pr8 virus infection may be attributed to a combination of factors, including cell death and potential limitations on the ability of the virus to continually infect new cells. Notably, our confocal images demonstrated a higher proportion of infected cells in the case of the Pr8 virus infection as compared to RSV. (**Figures S13-S14**). Under the fluorescent microscope, the evaluation of infection positive areas provided additional insights into the dynamics of viral infection at 24 or 48 h time-points. At 48 h, the infection-positive areas for both viruses were consistently higher compared to 24 h indicating a progression of viral spread and replication. Comparing Pr8 and RSV infection, the infection positive area was notably more extensive in the case of Pr8 virus infection. With ventilation, we observed a wider spread and higher infection rate. This increase in the infection rate correlated with both the amount of virus inoculum and the duration of infection, indicating a dose- and time-dependent relationship. This observation aligns with the previously discussed trend, where Pr8 virus exhibited a higher viral infectivity compared to RSV. The fluorescence microscopy data, represented by the infection-positive area, reinforced the notion that Pr8 virus had a more robust impact on host cells, resulting in a greater extent of infection-related changes in cell culture. Also, the MTT assay results were closely paralleled to the findings from the flow cytometry and confocal image analysis, where the relative cell viability determined using the MTT assay exhibited a similar trend to the viral infection levels assessed by flow cytometry (**Figure S15**). This consistency between the three quantification assays strengthens the reliability of our observations and underscores the accuracy of our viral infection assessments.

**Figure 4:**
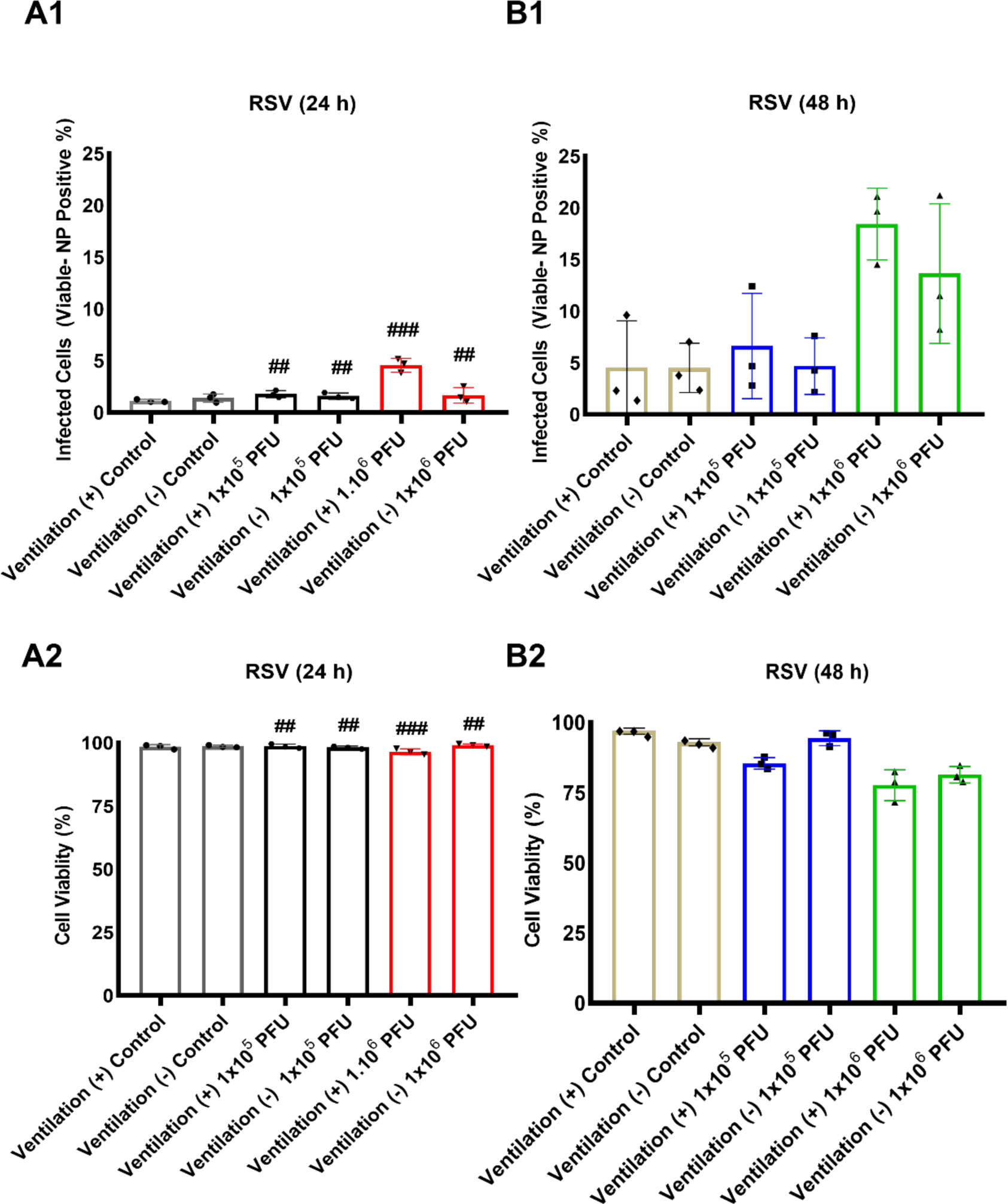
Quantification of RSV infection based on the Calcein and NP positive signal. **(A1 and B1)** Flow cytometry based on infected cells (%) at 24 or 48 h and **(A2 and B2)** cell viability (%) after 24 or 48 h of the viral infection (*n=*3; *p*<*0.05, *p**<*0.01 and *p***<*0.001; *p*^##^ <0.01 and *p*^###^ <0.001 indicate significance with respect to the control of the same group).

Finally, we compared the levels of pro-inflammatory cytokine and chemokine production in ALI cultures in response to the ventilation and respiratory viral infection. In uninfected samples, we found differences in protein concentrations between Ventilation (+)/Perfusion (+) and Ventilation (-)/Perfusion (+) groups for G-CSF, MCP-1/CCL2, and VEGF-A, while no significant difference was found for IL-6, IL-8, or CXCL10 (**Figures 5-6 and S16-S17**). For all significantly different protein expressions in the absence of infection, there was higher secretion in the Ventilation (+)/Perfusion (+) group, which suggests that ventilation status enhanced cytokine production. In the viral-infected cultures, ALI cultures infected by Pr8 had higher cytokine and chemokine concentrations irrespective of ventilation status compared to RSV, consistent with the higher rate of infection seen in influenza ALIs. In response to influenza infection at both 1×10^5^ and 1×10^6^ PFU, we found higher concentrations of G-CSF, IFN-β, IL-6, IL-8, and CXCL10 in Ventilation (-)/Perfusion (+) compared to Ventilation (+)/Perfusion (+), while MCP-1/CCL2 and VEGF-A had higher concentrations in the Ventilation (+)/Perfusion (+) group. A similar pattern was seen in RSV infected cultures for IL-8, but this was not replicated in other proteins measured. Compared to uninfected samples, many proteins were decreased compared to the uninfected control, which is likely related to differences in cell viability. However, there was very little CXCL10 produced at baseline, which significantly increased with infection and suggest its specificity. Overall, our data show that biologically and clinically relevant proteins can be measured in the supernatant of the Ventilation (+)/Perfusion (+) cultures, again showing the utility of this model system for studying respiratory viral infection.

**Figure 5:**
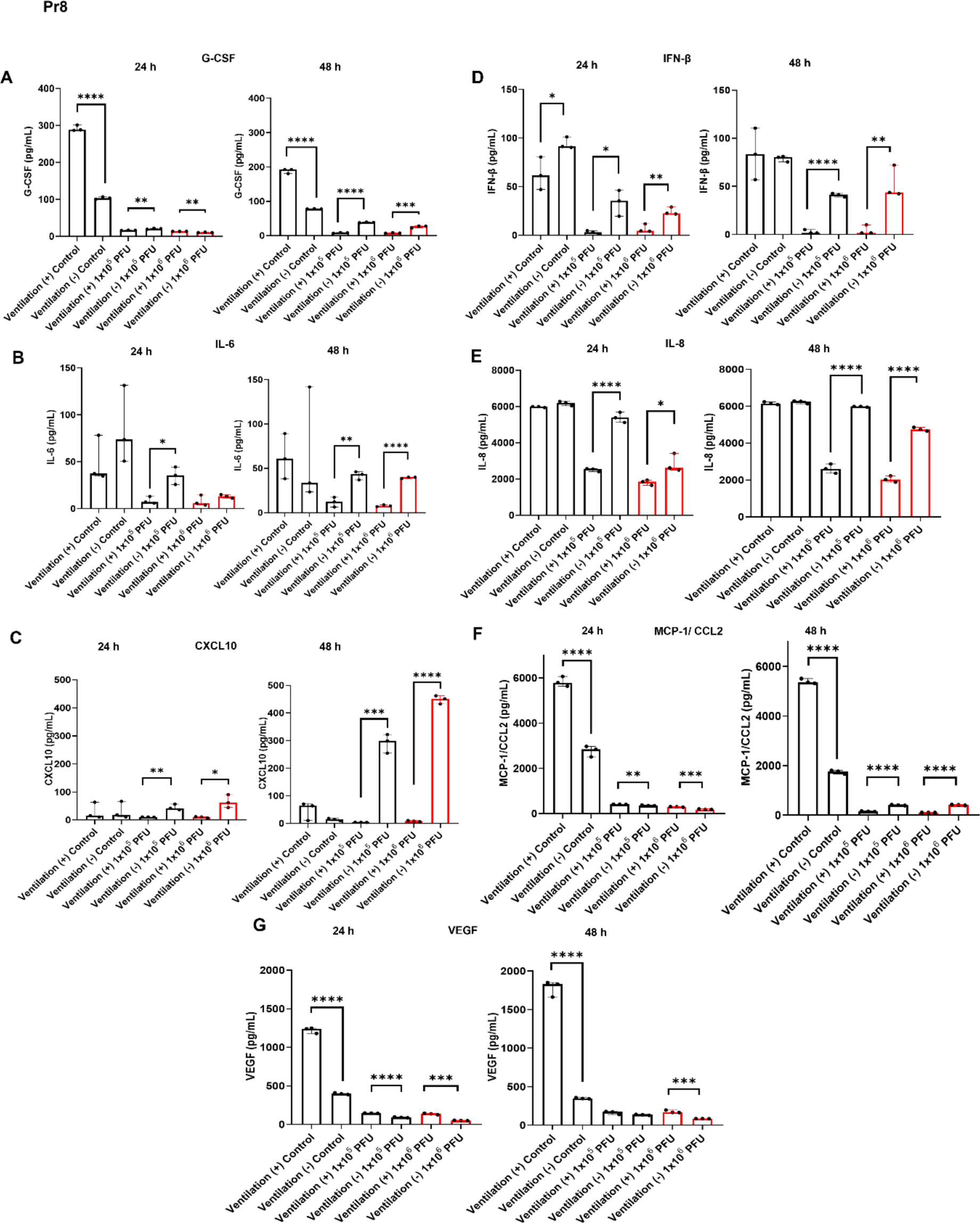
Lung device supernatant cytokines/chemokines measured using immunoassay across conditions. **(A)** G-CSF, **(B)** IL-6, **(C)** CXCL10, **(D)** IFN-β, **(E)** IL-8, **(F)** MCP-1/CCL2, and **(G)** VEGF proteins measured in supernatants under all conditions and at 24 or 48 h of Pr8 infection (*n*=3; *p*\*<0.05, *p*\*\*<0.01, *p*\*\*\*<0.001, *p*\*\*\*\*<0.0001).

**Figure 6:**
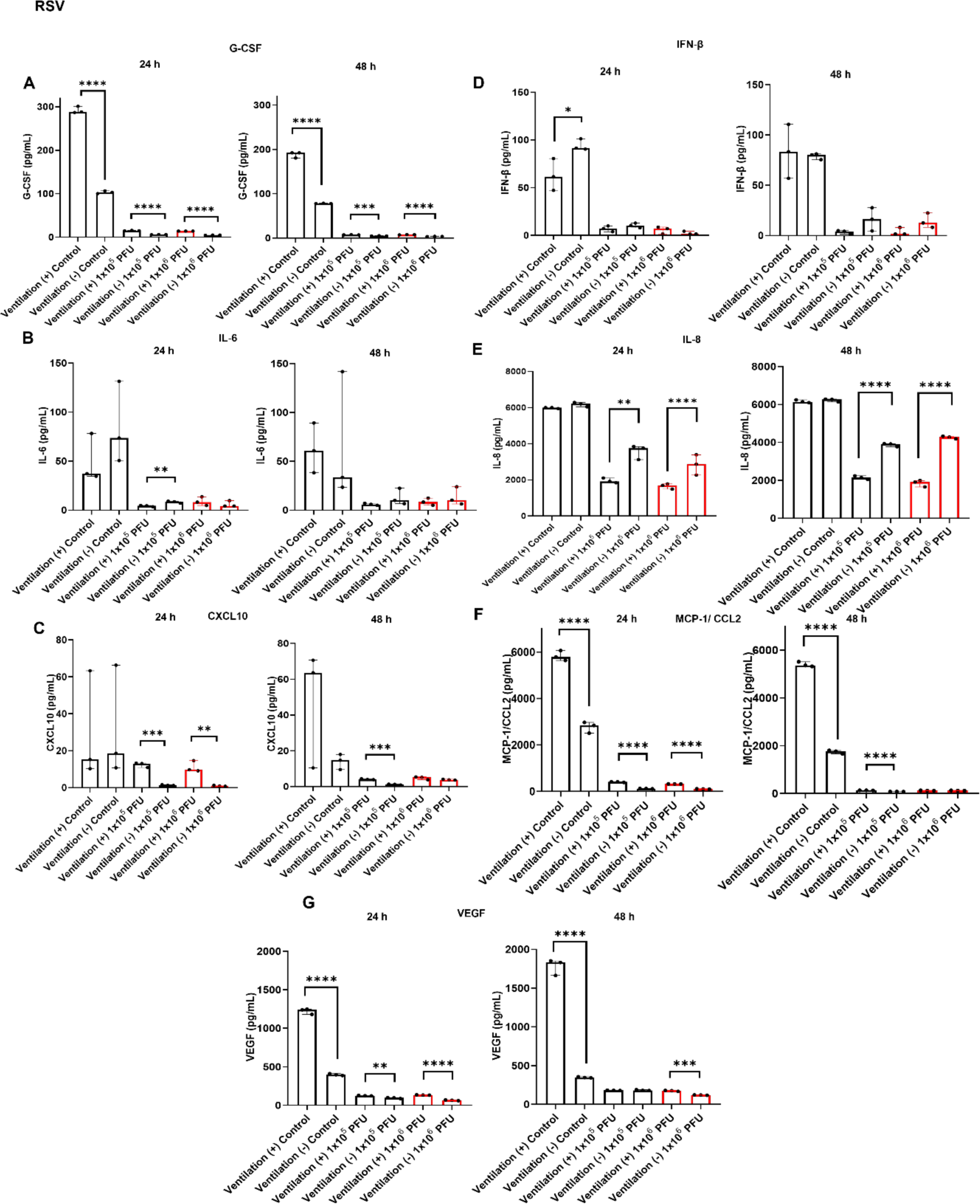
Lung device supernatant cytokines/chemokines measured using immunoassay across conditions. **(A)** G-CSF, **(B)** IL-6, **(C)** CXCL10**, (D)** IFN-β**, (E)** IL-8**, (F)** MCP-1/CCL2, and **(G)** VEGF proteins measured in supernatants under all conditions and at 24 or 48 h of RSV infection (*n*=3; *p*\*<0.05, *p*\*\*<0.01, *p*\*\*\*<0.001, *p*\*\*\*\*<0.0001).

## DISCUSSION AND OUTLOOK

To advance our knowledge of respiratory epithelial cells and their critical role in immune regulation and host defense, it is essential to study these cells in a setting that resembles their natural environment as closely as possible. Such a system could allow us to investigate differences among epithelial cells from individuals with respiratory diseases, the effects of infecting viruses and other lung pathogens and toxins (*49*, *50*). Conducting such research in a more contextually relevant manner, we may be able to shed light on the functions and interactions of respiratory epithelial cells and pave the way for targeted therapeutic interventions to improve respiratory outcomes.

The development of 3D printed lung models represents a significant advancement in the field of respiratory research. These models offer several advantages over traditional in-vitro and animal models (*51–53*), by providing a more physiologically-relevant platform for studying lung-specific immune responses and respiratory diseases. The application of 3D printing technology offers a promising avenue to create physiologically-relevant platforms by enabling the precise recreation of lung tissue microenvironments with human cell components, facilitating the investigation of immune interactions and disease mechanisms in a controlled and biologically accurate context. The ability to incorporate ventilation and perfusion components within a 3D architecture enables the simulation of complex cellular interactions and microenvironmental conditions found in human lungs. Undoubtedly, the ongoing COVID-19 pandemic has highlighted the significance of in-vitro models, including ALI cultures, in investigating the response to SARS-CoV-2 (*54*, *55*). These in-vitro models have played a crucial role in enabling researchers to study how the virus interacts with respiratory epithelial cells and the subsequent immune response (*13*, *56–58*). Such advancements in in-vitro platforms hold promise for accelerating our understanding of respiratory diseases, including viral infections, and can aid in the development of targeted therapies and preventive strategies.

In this study, we successfully developed a 3D printed ventilated and perfused human lung model that incorporated important the alveolar and vascular components. The use of 3D printing technology allowed for precise reconstruction of a model that resembles the human lung environment. To underscore the principal strengths of our lung model, a notable advantage is its capacity to facilitate mid-to high-throughput 3D printing. This capability empowered us to rapidly generate a substantial volume of lung tissue models for experiments. This enhanced throughput not only expedites the research process but also enables comprehensive investigations across a broader range of conditions and variables. As a result, our lung model’s ability to achieve mid-to high-throughput printing substantially accelerates the pace of scientific discovery and expands the scope of insights gained from our study. One of the key strengths of the ventilated model was its notable capability to facilitate the establishment and maintenance of tight junctions between lung epithelial cells, as evidenced by the presence of ZO-1, which was not detected in A549 cell cultures in previous publications (*59–61*). Ventilation, with its cyclic inflow and outflow of air, played a pivotal role in enhancing the formation of these tight junctions and promoting their robustness over time (*62*, *63*). Ventilation-induced mechanical forces effectively simulated the dynamic nature of the breathing process in the human lung. These mechanical cues are known to influence cellular behaviors and communication (*64*, *65*). In the context of lung epithelium, our study aimed to gain insights into the mechanical behavior of the 3D lung model under dynamic breathing conditions. Specifically, we investigated how mechanical stretch and shear stress resulting from ventilation influenced various aspects of cellular behavior and the overall performance of the lung epithelial barrier. One key observation was the greater displacement experienced by the sides of the alveolar lung cavity. This finding suggests that the mechanical stretch generated by the breathing process is particularly pronounced at the lateral aspects of the lung structure (*66*, *67*). This mechanical stretch is known to have significant implications for cellular behavior. Importantly, the mechanical cues associated with ventilation play a pivotal role in aligning and organizing lung epithelial cells. Consequently, the mechanical cues associated with ventilation likely contribute to the improved assembly and alignment of proteins, like ZO-1, which are pivotal for the formation of tight junctions (*68–71*). On the other hand, the displacement field analysis clearly indicates that the sides of the alveolar lung cavity experienced greater displacement in response to dynamic breathing conditions. This finding is significant as it suggests that the alveolar walls on the sides of the lung are more flexible or responsive to mechanical forces associated with breathing. Conversely, when examining the strain distribution, it becomes evident that the bottom of the lung experienced a notably higher level of strain compared to the rest of the structure. This unexpected result contradicts the displacement findings and warrants further investigation. This discrepancy suggests a complex interplay between tissue stiffness, mechanical constraints, and deformation mechanisms within the lung model. Comparisons with the native human lung’s mechanical behavior provide valuable context for interpreting these strain findings. The distal human lung is characterized by a network of interconnected alveoli surrounded by elastic fibers and connective tissue. During normal breathing, lungs experience varying degrees of strain, primarily attributed to the stretching and relaxation of their elastin-rich components. Studies on the human lung have shown that strain distribution is not uniform, with the basal regions of the lung experiencing higher strains due to gravitational effects and the influence of the surrounding structures (*46*, *47*) Strain distribution observed in the 3D lung model may be influenced by factors unique to the experimental setup. The differences in material properties, boundary conditions, and the absence of natural physiological processes, such as bronchial dilation (*72–74*), could contribute to variations between the model and the native lung. Additionally, the absence of blood flow and the lack of interaction with the thoracic cage in the model might affect strain patterns (*75–77*).

Mechanical stretch can also modulate cytokine or growth factor secretion as seen by the changes in secreted proteins seen in our assays (*78*). The consistent mechanical cues provided by ventilation contribute to the sustained functionality of the epithelial barrier, mimicking the lung’s physiological responses more faithfully. These findings highlight the importance of integrating physiological stimuli into in-vitro models, as they can significantly contribute to the model’s ability to recapitulate complex biological processes and behaviors. TEER and permeability studies collectively validated the successful formation of a functional epithelial barrier in the ventilated lung model, which is critical for simulating lung-specific immune responses and respiratory diseases accurately.

Lung infections, such as those caused by influenza viruses and RSV, are significant contributors to respiratory illnesses (*79*). These viruses exhibit similar symptoms and can result in severe illness primarily, for children and elderly populations (*80*, *81*). In the context of respiratory infections, it is noteworthy that influenza and RSV exhibit distinct preferences for targeting specific cells within the respiratory tract. Specifically, RSV infects ciliated cells in the bronchi and bronchioles (*82*), whereas human strains of influenza virus predominantly target non-ciliated cells (*83*). These varying cell tropisms arise from differences in the distribution of receptors responsible for viral attachment for each virus, which shape their infection patterns (*84*). To comprehensively assess the infection dynamics and cellular responses in our 3D printed lung model, we exposed our samples to Pr8 and RSV, at two virus concentrations (1×10^5^ and 1×10^6^ PFU) for two time periods (24 h and 48 h). In general, the infection rate increased in the 3D printed lung model ventilated groups. However, it is noteworthy that in the control groups, we observed that less than 2% of cells were infected with RSV. This discrepancy may be due to potential residual contaminants in the experimental setup. For instance, it is well known that soluble sulfated proteoglycans are highly inhibitory to RSV infection (*85*). When examining RSV infection, which is characterized by lower infection efficiency, we observed notably higher cell viability, mirroring the trend of reduced infection severity. This observation underscores the intricate relationship between ventilation and cell viability within our 3D printed lung model. As ventilation-induced mechanical forces influence the mechanical behavior of the lung model, it is plausible that high infection levels may render the cells more susceptible to the mechanical stresses associated with ventilation, contributing to reduced cell viability. Furthermore, we observed a lower rate of viability in the ventilated samples, which is likely due to the detrimental effects of the infection on the cells. This observation offers valuable insights into the underlying factors contributing to the relatively lower proportion of infected cells at the 48 h infection period.

The noticeable discrepancy in infection rates between the Pr8 and RSV within our 3D printed lung model warrants careful consideration. While our study revealed a higher infection rate with Pr8 virus, the underlying factors contributing to this disparity are not clear. One possibility lies in the different cellular receptor used by these viruses to initiate infection. Furthermore, the distribution and abundance of viral receptors on host cells might influence the infection pattern. Investigating these receptor profiles and their interaction with the viruses could shed light on why Pr8 exhibited a more potent infection rate in our model. An intriguing observation in our study was the temporal dynamics of infection rates, with a peak at 24 h that either stabilized or decreased at 48 h. This pattern prompts us to explore the mechanisms governing this phenomenon. One hypothesis is that host cells mount an initial antiviral response that limits viral spread. Alternatively, the viruses themselves may exhibit varying replication kinetics over time to elucidate this, further investigations into the interplay between host immune responses, viral replication dynamics, and differences in cell tropism for each virus are warranted (*86*) Understanding the reasons behind these temporal fluctuations in infection rates is essential for a comprehensive comprehension of viral behavior within our model. We observed that the ventilated samples showed higher infection compared to non-ventilated samples, which might seem counterintuitive since ventilation mimics physiological breathing conditions. This intriguing finding prompts us to critically evaluate the role of ventilation in viral infection dynamics. While ventilation is a natural lung defense mechanism, its impact on viral aerosolization and virus spread from infected to uninfected cells or enhanced viral entry into host cells should be explored further. A comprehensive examination of the interplay between ventilation, viral aerosolization, and cellular responses is crucial to fully grasp the implications of these findings. The significant decrease in cell viability observed in the ventilated group after 48 h of infection raises important questions about the underlying mechanisms. Understanding why the ventilated group experienced more significant cell death in response to infection compared to non-ventilated samples is pivotal. Ventilation introduces mechanical forces that could potentially exacerbate cellular stress or apoptosis in the presence of viral infection. Investigating the mechanistic links between ventilation, viral infection, and cellular viability could provide valuable insights into the physiological responses of lung tissue to respiratory viruses. Our findings underscore the intricate dynamics of viral infections within our lung model and warrant further exploration to uncover the underlying mechanisms influencing cell viability and infection progression. Furthermore, our observations highlight the model’s ability to accurately replicate infection dynamics similar to those observed in the human lung. We demonstrated the quantification of clinically and biologically significant proteins in the supernatant of ALI cultures from infected samples. The observation of decreased cytokine production in Ventilation (+) compared to Ventilation (-)cells was consistent with what has been reported in a prior study (*87*). In other studies, mechanical stretch, often at supraphysiologic levels meant to mimic mechanical ventilation have shown induction of cytokine responses (*78*). In this study, we noted high basal levels of cytokine production. However, influenza virus typically induced higher pro-inflammatory cytokine production compared to RSV consistent with the decreased infectivity rate of RSV. Interestingly, CXCL10, a known interferon induced chemokine was low in uninfected ALI but significantly increased with influenza infection in our study. CXCL10 has been shown to be a measure of disease severity in COVID-19 and may be a good marker of epithelial viral response in these ALI models (*88–90*). The discernible differences in infection susceptibility between the two viruses validate the model’s relevance and effectiveness in simulating viral infections in a manner that mimics physiological conditions (*91*, *92*).

By introducing A549 cells into our model, we aim to capture a broader spectrum of cellular interactions and complexities that are present within the human lung. Utilizing A549 cells was driven by several reasons that align with our research objectives. Firstly, A549 cells possess characteristics akin to epithelial cells found in the lung, enabling us to simulate certain lung tissue properties effectively. Secondly, these cells are readily available and have been extensively studied (*93*, *94*), ensuring accessibility and a wealth of existing knowledge to build upon. Thirdly, A549 cells exhibit responses to various stimuli that mirror aspects of lung behavior, providing a practical avenue for investigating cellular interactions (*95–97*). Nevertheless, it’s crucial to recognize the limitations of A549 cells compared to primary or induced pluripotent stem cell (iPSC)-derived alveolar cells. Primary cells, being directly derived from living organisms, offer a high degree of biological relevance by faithfully replicating the behavior and characteristics of human lung cells, making them ideal for physiologically accurate research. Their retention of donor-specific genetics is particularly valuable for personalized medicine studies and understanding individual responses to treatments and infections (*98*, *99*). However, working with primary alveolar epithelial cells presents challenges, as they can exhibit significant variability in behavior between different donors, potentially introducing experimental variability. Additionally, isolating progenitor cells requires selection of Type 2 alveolar cells from fresh lung tissue, complex medium, and three weeks to differentiate into columnar epithelium. Moreover, primary cells have a finite lifespan, necessitating a continuous supply of fresh cells for ongoing research endeavors (*100*, *101*). In contrast, iPSC-derived cells offer a more consistent source for alveolar cells and can be derived from blood or skin cells of different individuals, enabling their potential use for personalized medicine and patient-specific research (*102–104*). These cells can be guided to differentiate into diverse lung cell types, offering a broader perspective on lung complexity. Furthermore, our study unveiled unexpected characteristics in the A549 cells used, representing a notable departure from the typical attributes of normal human alveolar epithelial cells. Conventionally, alveolar cells do not exhibit mucin expression or possess ciliary structures, features that are customarily associated with airway cells. These unanticipated observations are particularly germane within the broader context of airway biology and underscore the need to revisit conventional assumptions about the cellular properties of alveolar cells, as they may have functional implications related to mucin expression and ciliary structures. However, they come with challenges related to differentiation protocols, time investment, and ethical considerations (*105–107*). In our study, the choice of A549 cells aligned with our research goals and the practical requirements of capturing essential aspects of lung biology while recognizing the potential advantages of iPSC-derived cells in other contexts.

This heightened fidelity in mimicking lung physiology and immune responses establishes our model as a valuable tool for in-depth investigations into lung ailments like chronic obstructive pulmonary disease, asthma, and respiratory viral infections. However, it is important to acknowledge that the 3D printed lung model possesses certain limitations that warrant addressing. While the model utilizes A549 cells, it may not fully encapsulate the diverse cell heterogeneity found in the human lung. Specifically, the model primarily focuses on respiratory epithelial cells and their interactions but may not comprehensively represent other crucial cell types present in the lung, such as airway cells, immune cells (e.g., macrophages, T cells), endothelial cells, and lung fibroblasts, which collectively contribute to the intricate cellular composition and functionality of the lung. For this reason, future efforts will concentrate on refining and expanding the range of cell types within the model to replicate the complexity of the lung microenvironment more authentically. Additionally, our study has primarily focused on establishing the lung tissue model’s structure, barrier function, and immediate response. To ensure the model’s reproducibility and reliability for extended experiments, it is crucial to conduct long-term culture and stability studies. Furthermore, the inclusion of additional components such as an endothelial barrier and the perfusion of immune cells would provide a more comprehensive and physiologically-relevant representation of the lung’s intricate interactions.

In conclusion, our innovative 3D printed human lung tissue model, featuring ventilation and perfusion capabilities, marks a significant advancement in exploring lung-specific respiratory infections. The model’s capacity to emulate the lung microenvironment with an established functional epithelial barrier positions it as a valuable asset for investigating a spectrum of lung biology aspects, host defense mechanisms against infections, and the formulation of targeted therapeutic strategies for lung-related conditions. As we continue refining and validating this 3D printed lung tissue model, it holds promise for furthering our comprehension of respiratory health and diseases, ultimately contributing to the enhancement of patient care and treatment approaches.

## MATERIALS AND METHODS

### Preparation of the gel and the fugitive ink

Before dissolving in Dulbecco’s Phosphate Buffered Saline (DPBS, 1x; Corning), gelatin powder (type A; 300 bloom from porcine skin, Sigma-Aldrich) was sterilized using Ultraviolet (UV) light for 3 h. 10% (weight (w)/volume(v)) gelatin was dissolved in DPBS, which was stirred at 90 °C for 2 h. Gelatin was then aliquoted and stored at 4 °C for further use. 20% (w/v) transglutaminase (TG; Moo Gloo TI Transglutaminase) was prepared by dissolving TG in DPBS, incubated at 37 °C for 1 h, and sterilized via filtering before use. 10% (w/v) gelatin was mixed with TG at 60 °C to prepare a 0.3% (w/v) TG immediately before printing.

The fugitive ink formulation was composed of two main components: 34, 38 or 42 % (w/v) Pluronic F-127 (Sigma-Aldrich) with varying concentrations of high molecular weight polyethylene oxide (PEO; average molecular weight ∼900,000, Sigma-Aldrich), at 0, 1, 1.5, and 2% (w/v). Ink preparation was performed in a cold room at 4 °C to ensure stability during the dissolution process. The fugitive ink was prepared by dissolving Pluronic F-127 and the specified concentrations of PEO in PBS. This preparation was carried out on a magnetic stirrer overnight to ensure thorough mixing and dissolution of the components. The controlled environment and extended mixing duration contributed to the stability and uniformity of the ink, which was crucial for its use in 3D printing.

### Rheological analysis and Compression testing

The rheological behavior of inks (with 34, 38 and 42% Pluronic, and 38% Pluronic with 1.5% PEO) were measured using an MCR 302 rheometer (Anton Paar), which was equipped with a 25-mm parallel-plate at room temperature. The shear rate sweep test was performed within a range of 0.1 to 100 s^-1^ shear rate. The frequency sweep test was performed within a range of 0.1 to 100 rad s^-1^ angular frequency. The amplitude sweep test was performed within a range of 0.01 to 100% shear strain. The thixotropic recovery sweep test was performed by applying 0.1 or 100 s^-1^ shear rates with 60 s intervals. Except for the frequency sweep test, all other tests were performed at a frequency of 1 Hz. Additionally, storage (G’) and loss modulus (G") and complex viscosity (|η*|) were obtained from frequency sweep tests performed at a frequency range of 0.1–100 Hz at a constant strain of 0.1%.

Unconfined compression tests were performed using a 5966 series advanced electromechanical testing system with a 10 kN load cell (Instron) according to the ASTM standard D395–18. Briefly, rectangular specimens 6 × 13 mm (height x diameter) were compressed at a rate of 1.3 mm/min to failure. Values were converted to stress-strain and the initial bulk modulus (kPa) was calculated from the initial gradient of the resulting curve (0–10% compressive strain) as shown in **Figure S2C**.

### Printability test

An Inkredible+ bioprinter (Cellink) was used to perform all 3D printing experiments at room temperature. Fugitive inks were transferred into syringe barrels with a plastic 22 GA (410 µm) tapered dispensing tip(s) (Taizhou Wuwu). A pressure control unit containing a built-in air pump was used to control the extrusion pressure. Printability was measured and analyzed for four inks (38% Pluronic with 0, 1, 1.5 or 2% (w/v) PEO). The distance between the nozzle tip and print-bed was set to be equal to the filament width (error < ±0.1 mm) in all experiments. To replicate the conditions of lung model fabrication, a layer of 10% (w/v) gelatin and 0.3 % (w/v) TG was cast on Petri dishes. Five minutes after casting, fugitive ink was printed on the gelatin surface in a bi-layer grid arrangement with varying pore sizes (e.g., 1 x 1, 2 x 2, 3 x 3, 4 x 4, and 5 x 5 mm x mm). The printed grid images of were obtained with a Nikon D810 camera with Nikon 105 micro lens attached and analyzed using ImageJ [National Institutes of Health (NIH)]. Equation (1) was used to calculate printability (*Pr*), where *Pr* = 1 indicated perfect printability of a square pore, Pr < 1 indicated a circular pore and Pr > 1 indicated an irregular-shaped pore (*35*).

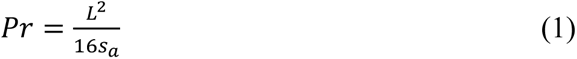

In Equation (1), *s_a_* was the actual area and *L* was the perimeter of the printed pores.

### Lung Model Fabrication

To fabricate the lung model, a custom-made device was prepared as shown in **Figure 1A**. The exterior of the device was printed with polylactic acid (PLA) using a 3D printer (X-Max, QIDI Tech). The side walls were laser cut from an acrylic sheet and PLA was glued to it. To image the lung model in this device, the bottom part was initially left open, and then a glass slide cover was glued to the bottom to seal it. Before fabricating the lung model, the device was sterilized with ethanol (70%) and UV light overnight. After sterilization, the first layer containing 10% (w/v) gelatin and 0.3% (w/v) TG was cast into the device. 5 min after casting, the gelatin layer was crosslinked and solidified. After solidification, a fugitive ink [38% (w/v) Pluronic with 1.5% (w/v) PEO] was printed on the gelatin surface using the Inkcredible+ bioprinter to obtain a perfusion channel (∼0.6 mm in diameter, ∼8 mm in length). Subsequently, a second layer of gelatin-TG mixture was deposited on the fugitive ink filaments to enclose them and allowed to solidify. After waiting another 5 min, fugitive ink was used to print the alveolar chamber (∼6 mm in diameter, ∼10 mm in height) followed by printing an airway (∼0.55 mm in diameter, ∼4 mm in height) on the second layer. Afterwards, the final layer of gelatin-TG mixture was cast to enclose the whole fugitive structure. After fabrication, the device was placed in an incubator for 10 min to facilitate full crosslinking between layers, and then placed in 4 °C for at least 4 h to liquify the fugitive ink. After liquefaction, the fugitive ink was flushed out with 4 ℃DPBS to open the perfusion channel, the alveolar chamber, and the airway.

### Lung Model Ventilation and Perfusion

Upon fabrication, the lung model was connected to a custom-made ventilation system. The ventilation system consisted of an air pump (Ultimus I, Nordson) to supply the air and vacuum, a humidification chamber to humidify the applied air, a pressure sensor to measure the pressure, and a syringe filter to sterilize the applied air (**Figure 1B)**. Humidifying the air was essential in our system to mimic the natural conditions of inhaled air, which also to maintain the hydration of the gelatin-based device, which ensures the structural stability of the respiratory model. In other words, this process facilitated the replication of moisture levels present in human lungs and safeguarded the integrity of the lung model. The pressure sensor was placed upstream of the lung model to measure the applied air pressure because space limitations prevented the sensor from being placed inside the chamber. The air pressure and vacuum were set to obtain ±1-1.5 cm H_2_O at the pressure sensor. After humidified air was applied continuously with a 3 -sec inhalation and 3 -sec exhalation cycle, the lung model was connected to the perfusion pump.

### Cell culture

For cell culture experiments, the fabricated lung model was prepared as described in the previous section. Cell seeding took place after the fugitive ink was flushed out of the perfusion channel and the alveolar chamber and the airway were opened. To maintain hydration of the lung model, additional cell culture media was carefully introduced to the exterior sides of the model as shown in **Figure 1A**. A549 Human adenocarcinoma cells (ATCC# CCL-185) obtained from the American Type Culture Collection (ATCC) were grown in F-12K Medium (Kaighn’s Modification of Ham’s F-12 Medium, ATCC). The medium was supplemented with 10% fetal bovine serum (FBS, R&D Systems), 1% Penicillin/Streptomycin (Corning) and 1% sodium pyruvate (Corning). Cells were maintained in a humidified incubator at 37 °C with 5% CO_2_. To generate ALI cultures, the apical surface of alveolar chambers was coated with 1 % (w/v) collagen type IV (Sigma-Aldrich), 0.5×10^6^ A549 cells were then seeded onto the coated chambers and cultured for 10 days. The culture system utilized a mixture of PneumaCult-ALI basal medium [supplemented with PneumaCult-ALI supplement (10x), PneumaCult-ALI Maintenance Supplement (100x), Hydrocortisone stock solution, Heparin solution, and Rho-kinase inhibitor (Rock Inhibitor)], all sourced from Stem Cell Technologies. During the initial culture phase, from seeding day to the start of ventilation, samples were fully submerged in the medium to optimize cell growth.

Subsequently, on Day 0 of ventilation, the apical culture medium was removed, and the basal chamber medium was replenished with PneumaCult ALI medium without Rock Inhibitor. This medium was replenished every other day until Day 10, which ensured consistent nutrient supply. A continuous perfusion setup was employed, wherein the cell medium was perfused at a flow rate of 1 ml/min for the first 24 h and increased to 2 ml/min for the subsequent days of culture until Day 10, unless otherwise specified. Four groups were studied to analyze the effects of ventilation and perfusion in the 3D printed models: Ventilation (-)/Perfusion (-), Ventilation (+)/Perfusion (-), Ventilation (-)/Perfusion (+), and Ventilation (+)/Perfusion (+). This approach was for a systematic examination of the roles of ventilation and perfusion individually and in combination, to determine their impact on the response of the lung model. For the Perfusion (-) groups, which did not involve continuous perfusion of medium through the lung model, we followed a modified protocol, introducing the medium by gently submerging the cultures in medium. This involved precise delivery of culture medium to the basal surfaces of the lung model, which ensures homogeneous distribution while minimizing any potential disturbance to cultured cells. The volume of medium introduced was consistent across all Perfusion (-) groups.

### Computational Modelling

The elastic deformation of the ventilated lung was modeled using COMSOL Multiphysics (COMSOL, Inc). We modeled the gel in the simulation as an incompressible hyper-elastic neo-Hookean material with an elastic modulus of 8.4 kPa, which was determined from the compression test (*36*). The side and bottom walls were prescribed fixed boundary conditions while the top surface was free to move. The surface normal pressure boundary condition was applied on the lung cavity with a maximum pressure of 1.1 mmHg. A stationary solver predicted the steady state shape of the deformed structure.

### Immunofluorescence staining

Immunofluorescent imaging was used to analyze the expression of ZO-1, α-tubulin and MUC5B. A549 cells were fixed overnight at 4 °C using 4% paraformaldehyde (PFA, Santa Cruz). After fixation, the cells were permeabilized with 0.1% Triton X-100 (Sigma-Aldrich) in DPBS for 10 min. Subsequently, a blocking solution consisting of 10% normal goat serum (Abcam), 1% bovine serum albumin (BSA, Research Products International), 0.3 M glycine (Sigma-Aldrich), and 0.1% Tween 20 (Sigma-Aldrich) in 10 mL DPBS was applied to the samples and incubated for 1 h at room temperature. Monoclonal antibodies specific to ZO-1 (Invitrogen), Alpha Tubulin (TUBA4A; Origene), and MUC5B (Invitrogen) were each diluted to a concentration of 5 mg/mL in the blocking solution. Samples were then incubated with these antibodies overnight at 4 °C. Subsequently, the samples were incubated with Alexa 647-conjugated fragment of goat anti-mouse IgG (Invitrogen) for 2 h at room temperature. Concurrently, cell nuclei were stained with a 4’,6-diamidino-2-phenylindole (DAPI) solution (1 mg/mL) for 2 h at room temperature. After each step, cells were carefully washed three times for a total of 3 min. The washing procedure ensured the removal of excess reagents and provided a clean sample for subsequent steps. Finally, stained samples were imaged using a confocal microscope (LSM880, Zeiss) at varying magnifications (e.g., 5x, 10x, 20x and 40x).

### Tight-junction analysis

To assess the tight junction density in different groups on Days 4, 7 and 10, the acquired ZO-1 images were subjected to semi-quantitative analysis using ImageJ Fiji software. A single line was manually drawn on the fluorescent images to extract the intensity profile. The peaks in the intensity profile were then identified. Subsequently, the position of each peak was converted to the corresponding scale in micrometers (µm), and the distance between two adjacent peaks was measured. This allowed for the quantification of the ZO-1-to-ZO-1 distance, providing an objective measure of tight-junction density.

### TEER measurements

TEER values were measured across the epithelial layer using an EVOM2 epithelial voltage meter (World Precision Instruments), equipped with a STX2/chopstick electrode. Electrodes were sterilized and calibrated in accordance with the manufacturer’s instructions. Cells were washed with pre-warmed DPBS before TEER measurements. Subsequently, pre-equilibrated ALI medium was added to both inside and outside of the device. To allow the monolayers to stabilize, the devices were incubated for approximately 5 min inside an incubator, ensuring a steady potential was reached. TEER values were then recorded using the voltage meter. To determine the sample resistance, the resistance of the acellular lung models was subtracted from the measured values. The resistance of each device, measured in ohms (Ω), was calculated using a previously described method (*37*).

### Diffusional permeability

To assess the diffusional permeability of the alveolar chamber, a solution containing 20 µg/ml of FITC-conjugated 4 kDa Dextran (Sigma Aldrich) was added into the alveolar chamber using the F12K medium. Images were captured at 5 min intervals using a fluorescence microscope (AxioObserver, Zeiss) for 1 day. Subsequently, the diffusion of Dextran and the resulting changes in fluorescence intensity were quantified using ImageJ. These measurements were conducted on samples with and without cells; three replicates (*n*=3) for each condition.

### Quantification of A549 markers

Following the acquisition of α-Tubulin and MUC5B images, fluorescently labeled areas were quantified using ImageJ Fiji. For the cilia area, a specific region, measuring 10^5^ mm^2^, was selected and analyzed to obtain the absolute value of the area. In contrast, for the MUC5B positive area, the measurement was represented as a percentage of the total area. This approach allowed for the quantification and comparison of the respective areas in a standardized manner.

### Quantitative reverse transcription polymerase chain reaction (qRT-PCR)

Cells were washed twice with PBS and then harvested with Trypsin and collected by centrifugation at 1,500 xg for 5 min and homogenized in TRIzol reagent (Life Technologies) at Day 10. PureLink RNA Mini Kit (ThermoFisher) was used to isolate total RNA from samples according to the manufacturer’s protocol. Nanodrop (Thermo Fisher Scientific) was used for measuring RNA concentration and reverse transcription was performed using AccuPower® CycleScript RT PreMix (BIONEER) following the manufacturer’s instructions. Gene expression was analyzed quantitatively with SYBR Green (Thermo Fisher Scientific) using a QuantStudio 3 PCR system (Thermo Fisher Scientific). Genes whose expression were quantified included aquaporin-5 (AQP-5), mucin-5B (MUC5B), mucin-5A (MUC5A), mucin-5AC (MUC5AC), surfactant protein-B (SFTPB), and surfactant protein-D (SFTPD), as listed in **Table 1**. All genes were normalized to the expression of the housekeeping β-actin, and 2^3ΔΔ56^was used to calculate the gene expression levels relative to the liquid-liquid culture at Day 1.

**Table 1.**
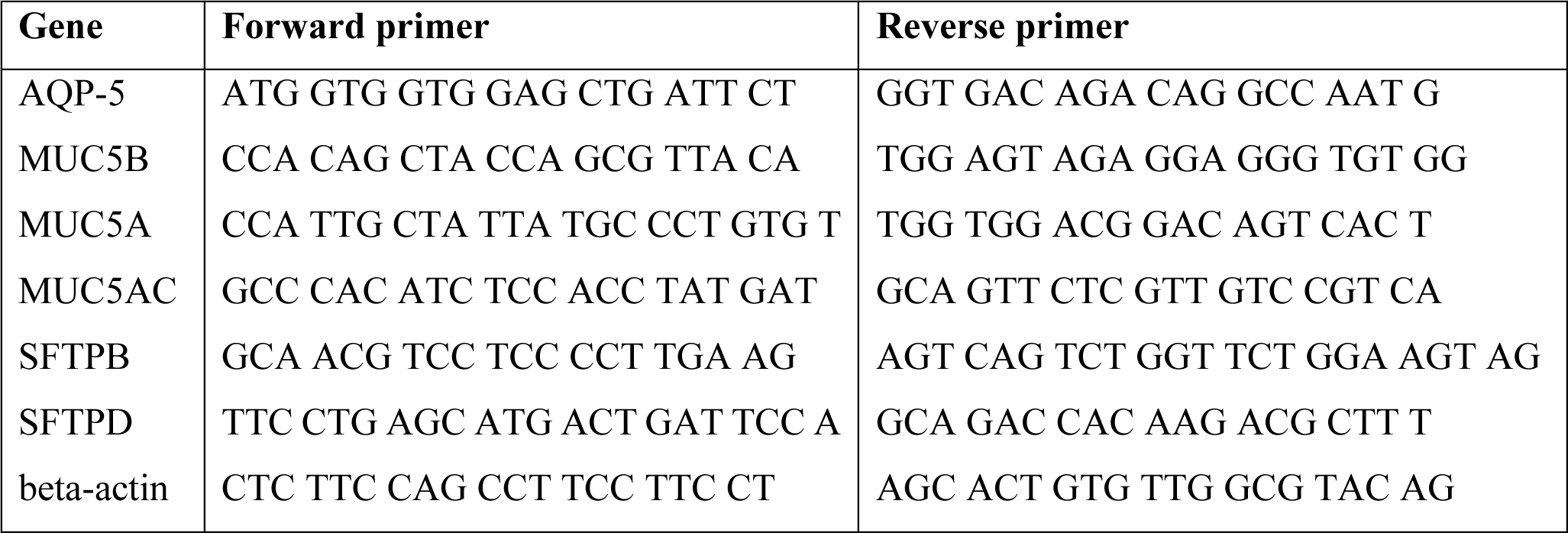
Primers used for qRT-PCR quantification.

### Viral Infection

Influenza virus (A/Puerto Rico/8/34 H1N1, abbreviated as Pr8), expressing mCherry fused to the viral non-structural 1 (NS1) protein was described before (*38*) and generously provided by Dr. Michael Schotsaert (Mount Sinai, NY). The respiratory syncytial virus (RSV), expressing the red fluorescent protein (RFP), was generously provided by Dr. Mark Peeples (Nationwide Children’s Hospital, OH). Aliquots of each virus were stored at -80 °C. Prior to infection, the apical surface of ALI cultures was washed 3 times with 1x PBS for 5 min at 37 °C. Subsequently, ALI samples were exposed to 100 µL of solution containing each virus on the apical side. The virus was diluted in serum-free Pneumacult ALI medium, which resulted in a final concentration of 1×10^5^ and 1×10^6^ PFU of Pr8-mCherry and RSV-RFP, respectively. After incubation for 24 or 48 h, 100 µL of PBS was added to the apical side of the ALI cultures for 15 min at 37 °C.

### Flow cytometry

To accurately segregate A549 cells from other cell types, a correlation analysis of both forward (indicates cell size) and side scatter (indicates cell granularity) was performed by flowcytometry (BD LSR-Fortessa) capturing a minimum of 5,000 cellular events in a forward-side scatter gate based on cell size and granularity. Data were analyzed using FlowJo v10.3 (BD Bioscience) to gate on single, live (Calcein violet) cells. We next gated on infected cells mCherry(PR8) or RFP(RSV). Flow cytometric monitoring of virus infection was analyzed using recently described studies (*39*, *40*).

### Cytotoxicity Assay

Cell viability was assessed using the methylthiazol tetrazolium (MTT, Sigma-Aldrich) assay to determine the viability of A549 cells after exposure to virus. MTT assay was conducted in 96-well plates with cells at 80-90% confluency (2 x 10^4^ cells per well) exposed to 1×10^5^ or 1×10^6^ PFU of RFP-RSV or mCherry-Pr8 viruses for 24 or 48 h. After exposure to the viral particles, 10 µL of MTT solution (5 mg/mL in PBS) was added to each well and incubated at 37 °C for 4 h. Formation of purple formazan crystals was assessed every 30 min. Once the proper formazan crystal formation was observed, the content in each well was completely aspirated and 100 µl of 100% dimethylsulfoxide (DMSO) was added to each well. The plate was mixed at room temperature for total of 4h, and the absorbance at 570 nm was measured using a microplate reader (Beckman Coulter AD 340C).

### Supernatant Analysis

Supernatants were collected at 24 or 48 h from the Ventilation (+)/ Perfusion (+) and Ventilation (-)/ Perfusion (+) groups of uninfected and viral-infected samples. For each condition, three experimental replicates were obtained. Cell-free culture supernatants were frozen at -80 °C and underwent two freeze-thaw cycles prior to analysis. Protein concentrations of granulocyte-colony stimulating factor (G-CSF), vascular endothelial growth factor (VEGF-A), interleukin-1 receptor antagonist (IL-1RA), Interferon-β (IFN-β), Interleukins-6, -8, -1β (IL-6, IL-8, IL-1β), and chemokines (CXCL-10, CCL-2, CCL3) were measured using a multiplex assay (U-PLEX Viral Combo 1, Meso Scale Diagnostics). The culture supernatant was diluted 1:3 based on preliminary dilution tests to ensure that the analyte would be within the detection range. The upper and lower limits of detection for each of the assay were comparable to the manufacturer’s standards. The upper and lower limits of detection of the analytes were: G-CSF (0.07-1930 pg/mL), IFN-β (2-106,000 pg/mL), IL-1β (0.03-4,830 pg/mL), IL-1RA (1.2-4720 pg/mL), IL-6 (0.18-1,930 pg/mL), IL-8 (0.04-2,470 pg/mL), CXCL10/IP-10 (0.3-11,100 pg/mL), MCP-1/CCL2/ MCP-1 (0.23-6,100 pg/mL), CCL3/MIP-1a/CCL3 (2.4-5,830 pg/mL), and VEGF-A (0.07-2,670 pg/mL). To avoid batch effect, samples were randomized and distributed equally across plates, and assays were run with the same reagents on the same day. If a sample measurement was below the lower limit of detection, the data were imputed using half of the value of the lower limit of detection. IFN-β had 11 imputed values, IL-1RA had 3 imputed values, and MIP-1a had 2 imputed values. We included assays that had <25% coefficient of variance between two plates, thus IL-1β, IL-1RA, and MIP-1a/CCL3 were removed from the final analysis.

### Statistical analysis

The collected data were analyzed and visualized using GraphPad Prism 8 software (GraphPad Software). Statistical analysis was performed using SPSS 28 software (SPSS, Inc.). To compare the differences among multiple groups, one-way analysis of variance (ANOVA) was conducted, followed by a post-hoc Tukey test. For analysis of cell culture supernatant protein measurements, values were compared using a two-tailed t-test. Significance levels were denoted as p*< 0.05, p** < 0.01, p*** < 0.001, and p**** < 0.0001 indicating the statistical significance of the observed differences.

## Acknowledgments

This research was primarily supported by National Institute of Allergy and Infectious Diseases Award U19AI142733 and in part by National Institute of Biomedical Imaging and Bioengineering Award R01EB034566 and 2236 CoCirculation2 of TUBITAK award 121C359. We thank Aaron O’Donnell, Vaibhav Pal, Shweta Saini and Beyza Albayrak from Penn State for their assistance with device preparations. The authors are also thankful to Ethan M. Gerhard and Dr. Jian Yang from Penn State for compression testing experiments.

## Author Contributions

I.D.D., D.B., and I.T.O developed the ideas and designed the experimental plan. I.D.D. performed cell culture, immunoimaging, TEER measurements, permeability experiments, viral infection, and all related data analysis, and led the manuscript writing. M.A.A. performed 3D printing and developed the ventilation system, and performed related analysis. D.N.K. assisted with the lung model development and related data analysis. S.E.H. and C.M. performed the cytokine expression experiments and related data analysis. M.N. assisted with perfusion, performed flow cytometry experiments and participated in the related data analysis. S.H.A.R. performed the simulation study. N.C. performed qRT-PCR and related data analysis. D.C.C. performed the data analysis for flow cytometry. W.P. and M.S. cultured Pr8 and P.C. and M.E.P. cultured RSV viruses, and they all provided critical insight into viral infection and related data analysis. K.P. and J.K. provided critical insights into viral infection and cytokine data analysis. All the authors contributed to the manuscript writing.

