## Supplementary Information for "A 3D Printed Ventilated Perfused Lung Model Platform to Dissect the Lung’s Response to Viral Infection in the Presence of Respiration"

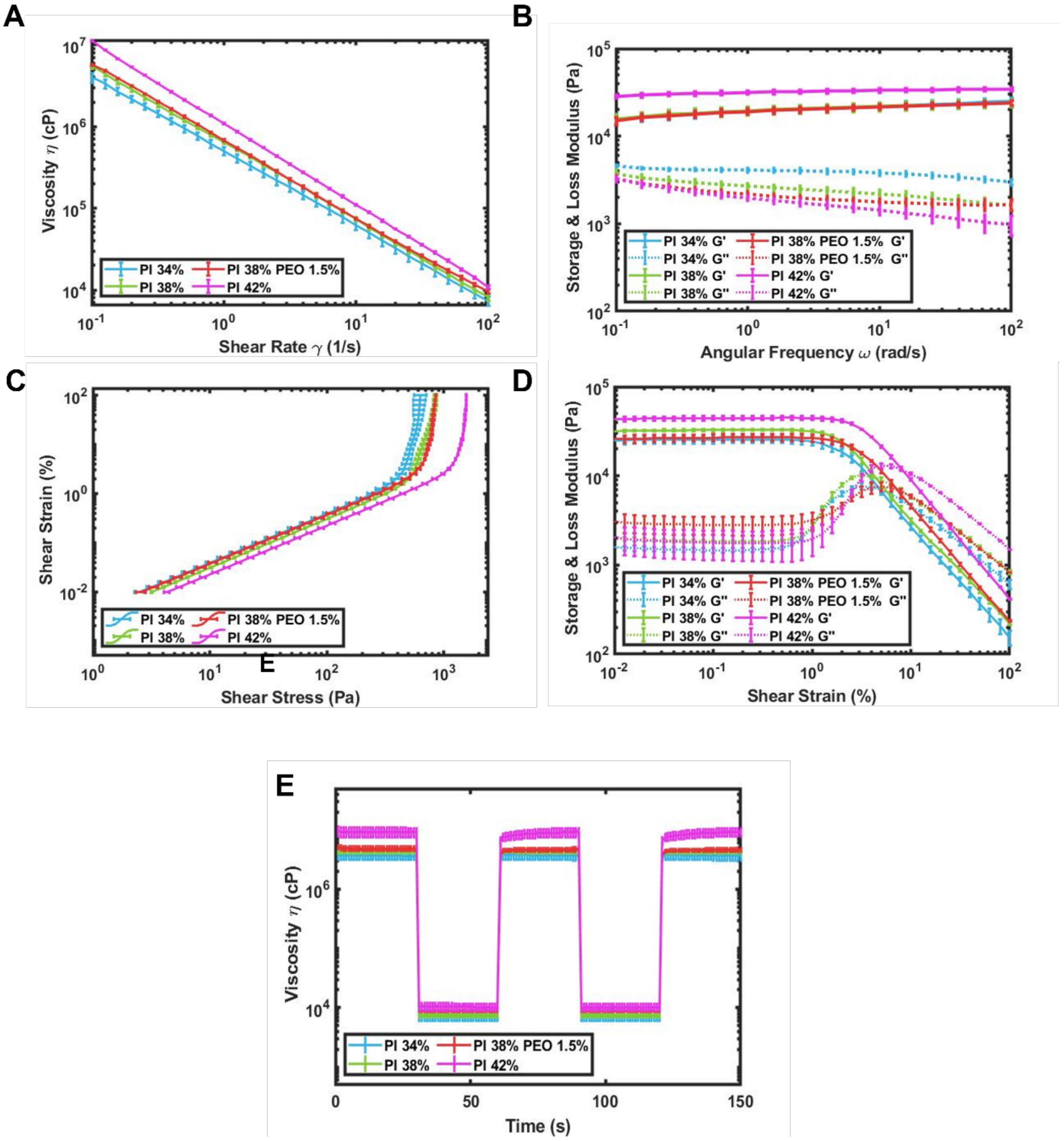


**Figure S1:** Rheological properties and printability of Pluronic F-127-based fugitive inks loaded with PEO. **(A)** Shear rate sweep test at a shear rate ranging from 0.1 to 100 $s^{-1}$, **(B)** frequency sweep test at a shear rate ranging from 0.1 to 100 rad $s^{-1}$, **(C)** shear strain vs. shear stress curves, **(D)** amplitude sweep test at a shear strain ranging from 0.01 to 100%, **(E)** and the self-recovery test of the fugitive inks. Viscosity measurements were performed at five intervals with alternating shear rates of 0.1 $s^{-1}$ for 30 s and 100 $s^{-1}$ for 30 s.

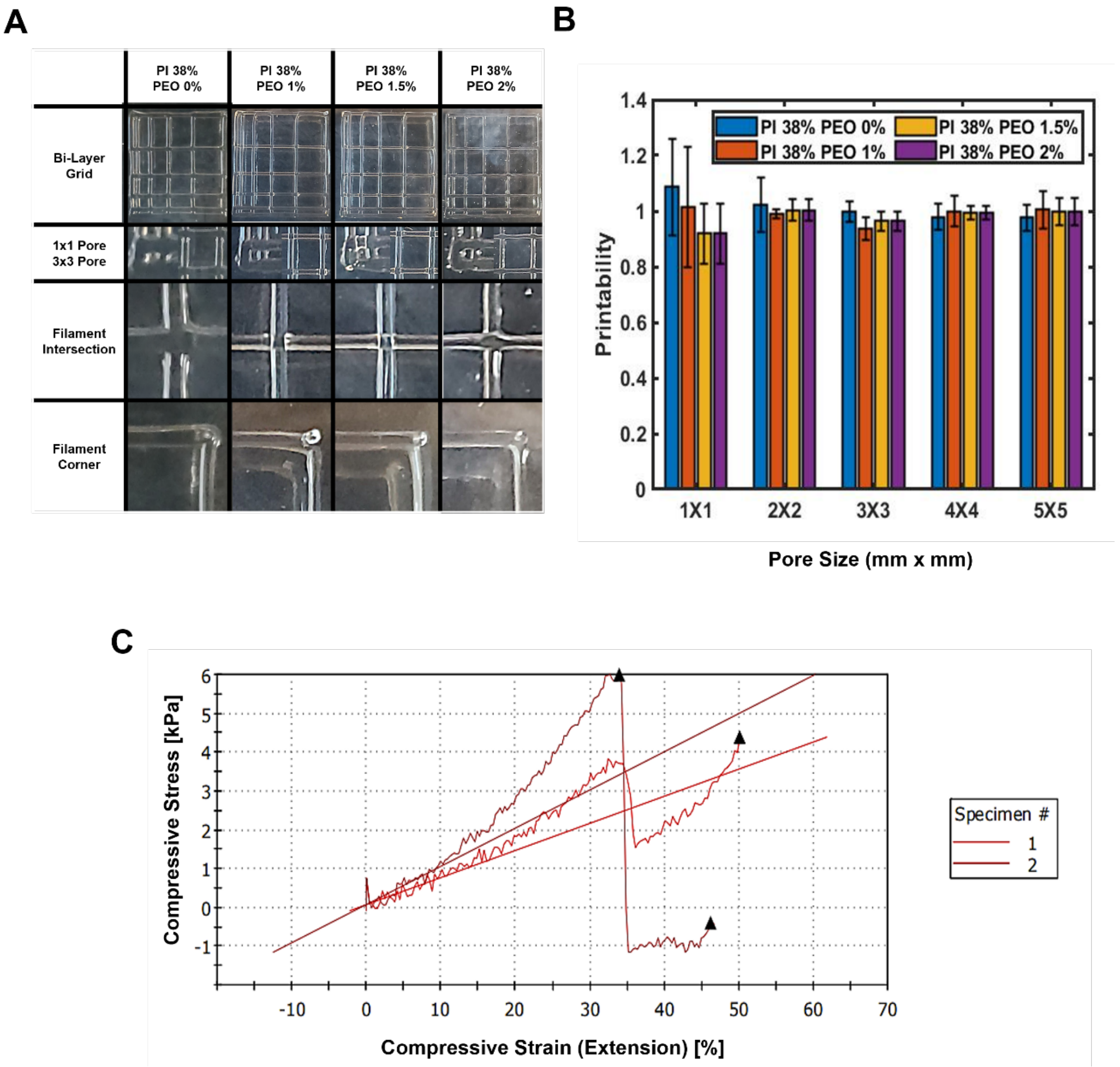


**Figure S2: (A)** 3D Printed bi-layer grid structures with increasing pore size on the crosslinked gelatin surface. **(B)** Printability of fugitive inks with different pore sizes (*n*=3). **(C)** Compression test results for distinct Gelatin+TG molds (*n*=2).

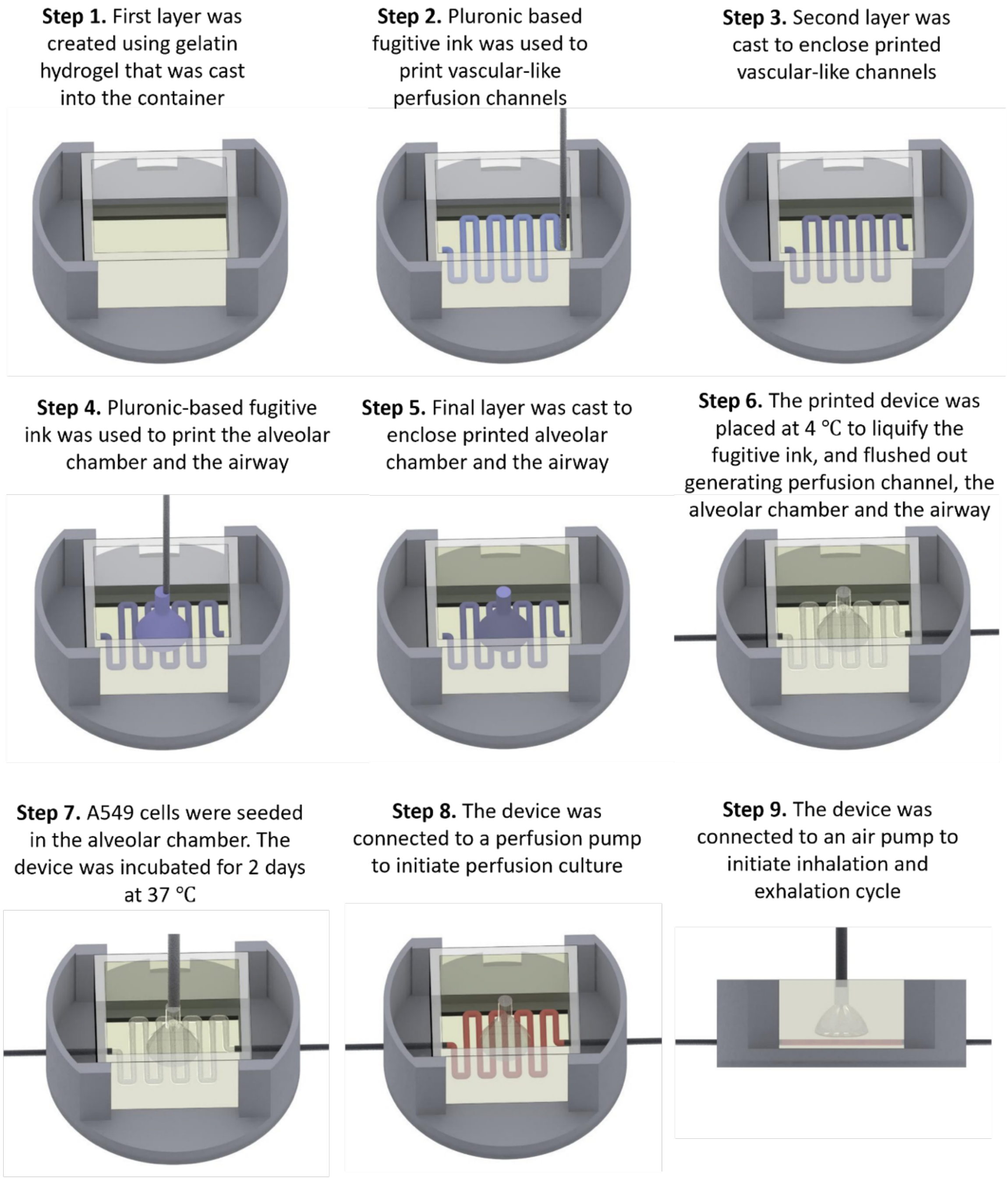


**Figure S3:** Detailed schematic demonstrating the step-by-step development of the lung model.

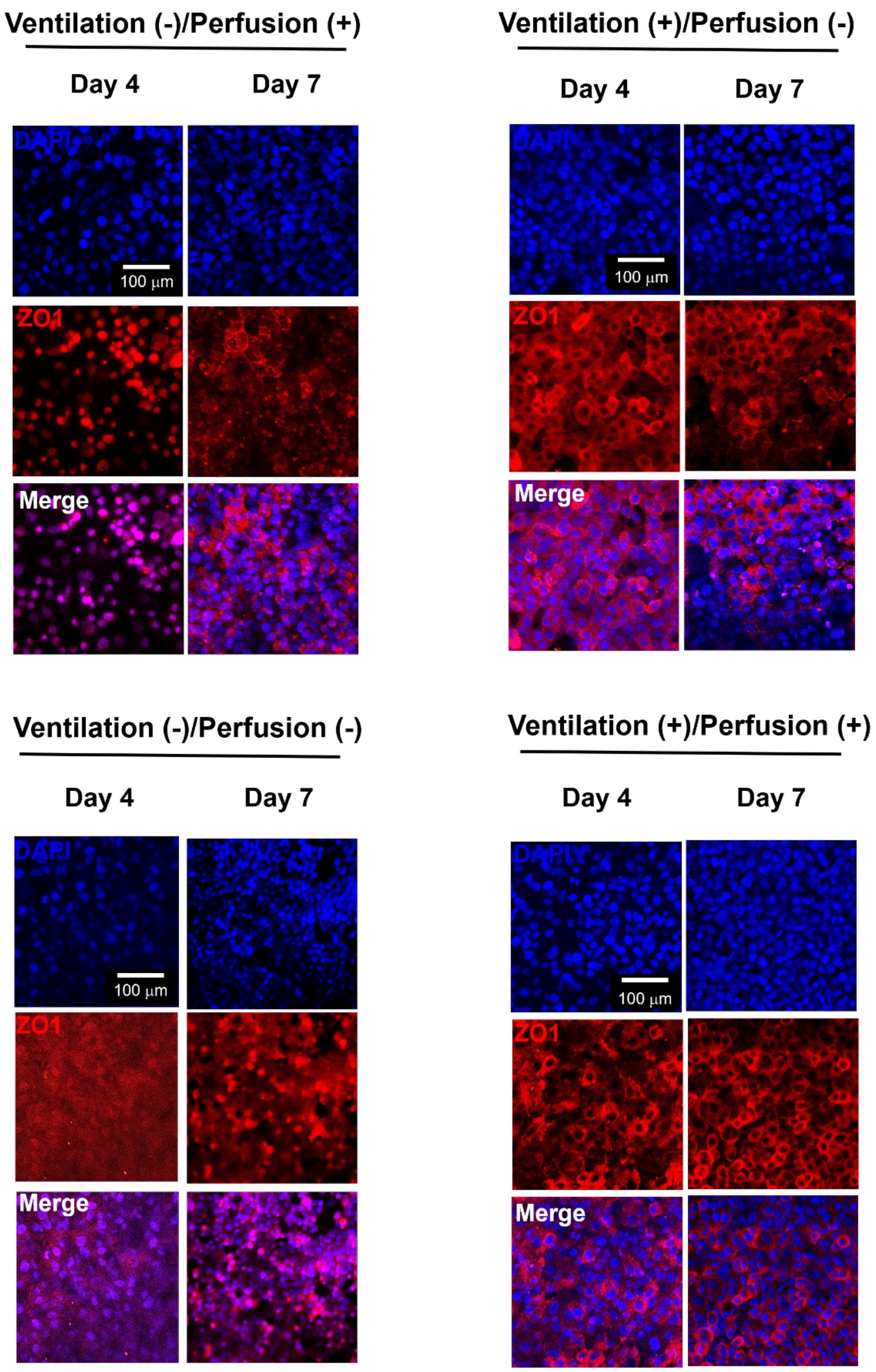


**Figure S4:** Evaluation of tight junctions in A549 cells using ZO-1 staining on Days 4 and 7.

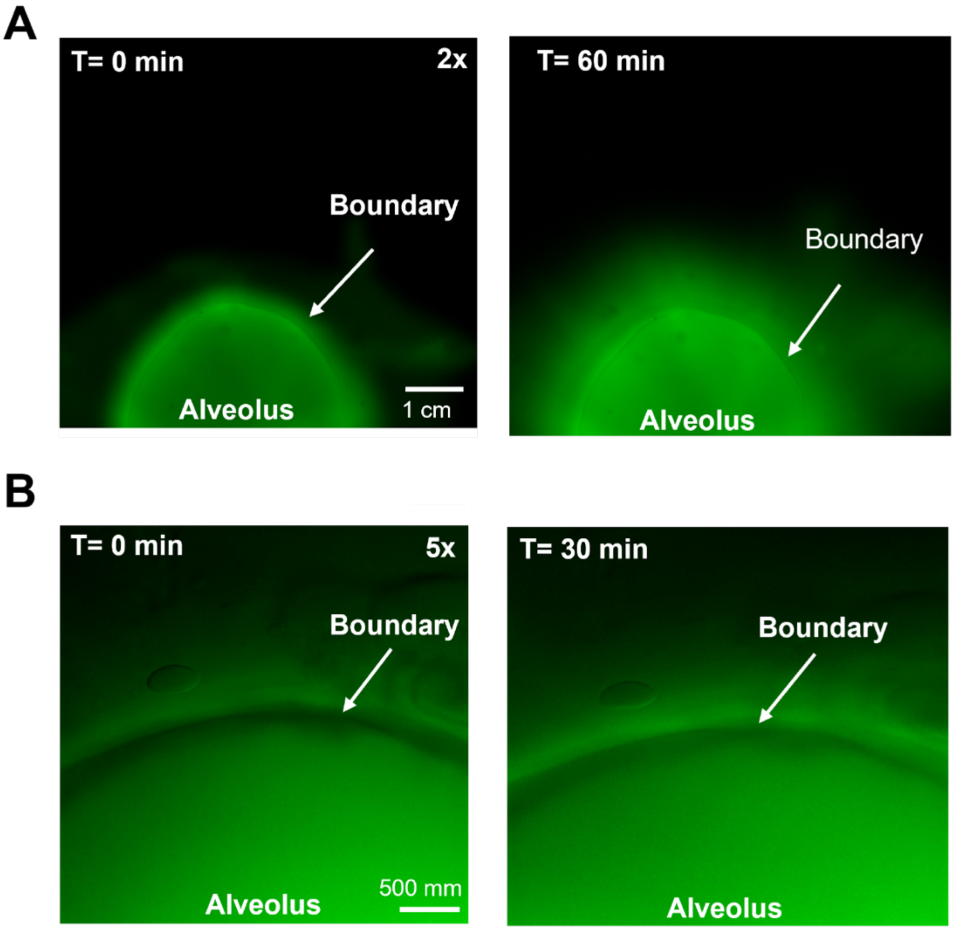


**Figure S5:** Representative images of Dextran diffusion for the acellular alveolar model **(A)** at T=0 and T=60 min (2x), **(B)** at T=0 and T=60 min (5x).

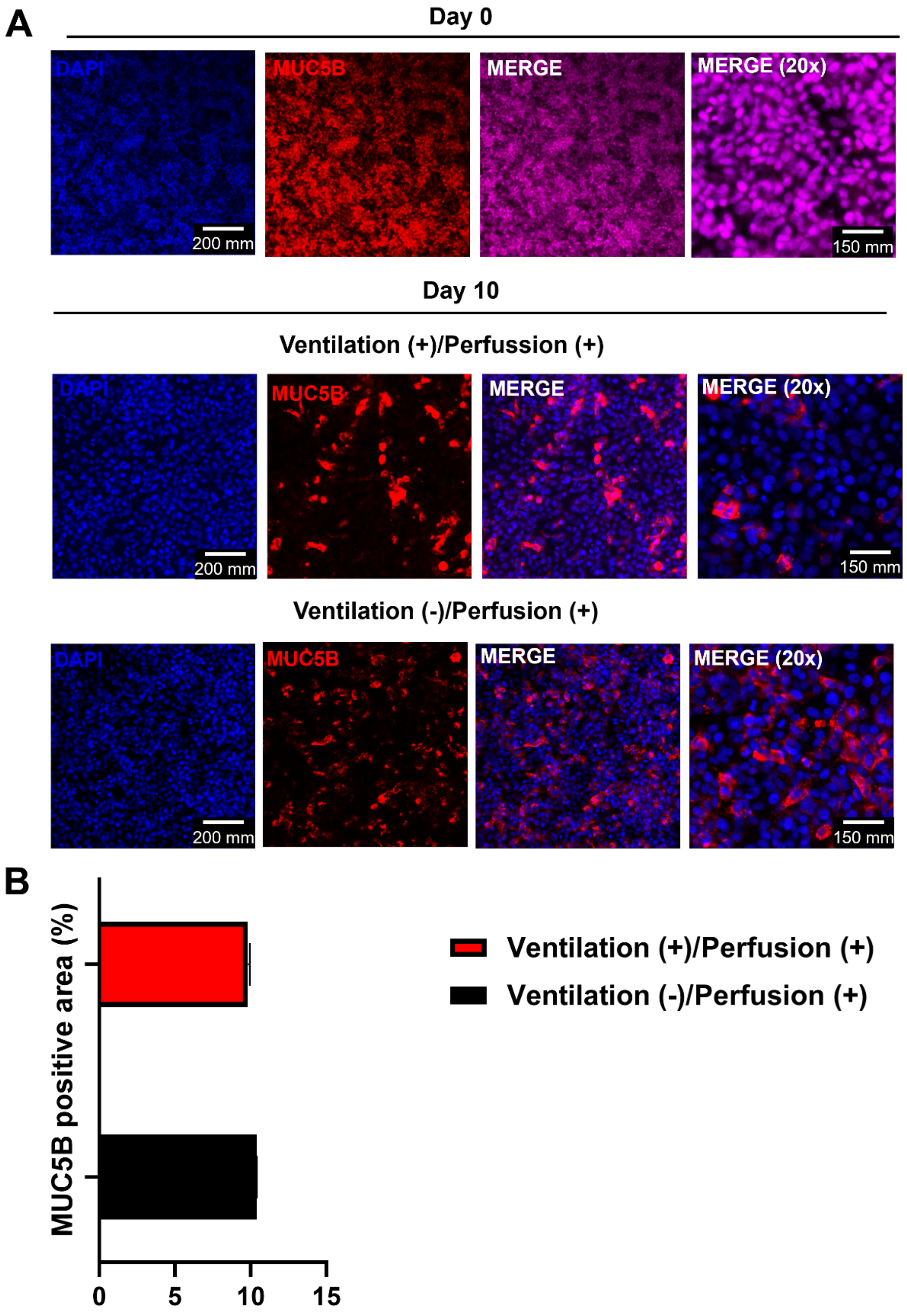


**Figure S6:** Representative immunofluorescent images of A549 cells characterized for MUC5B (red) and DAPI (blue) staining.

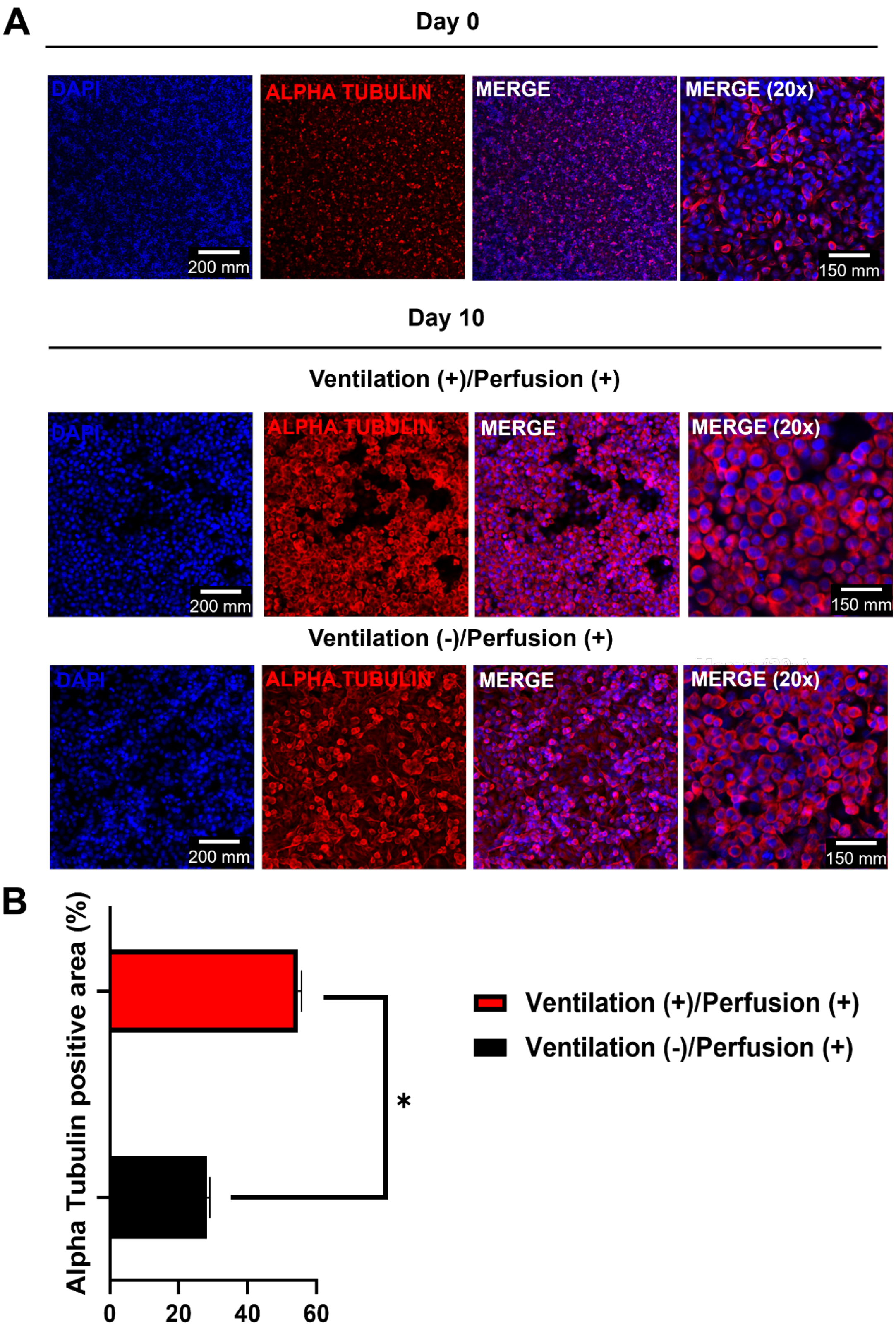


**Figure S7:** Representative immunofluorescent images of A549 cells characterized for acetylated α-tubulin (red) and DAPI (blue) staining ($n$=3; $p^{*}$<0.05).

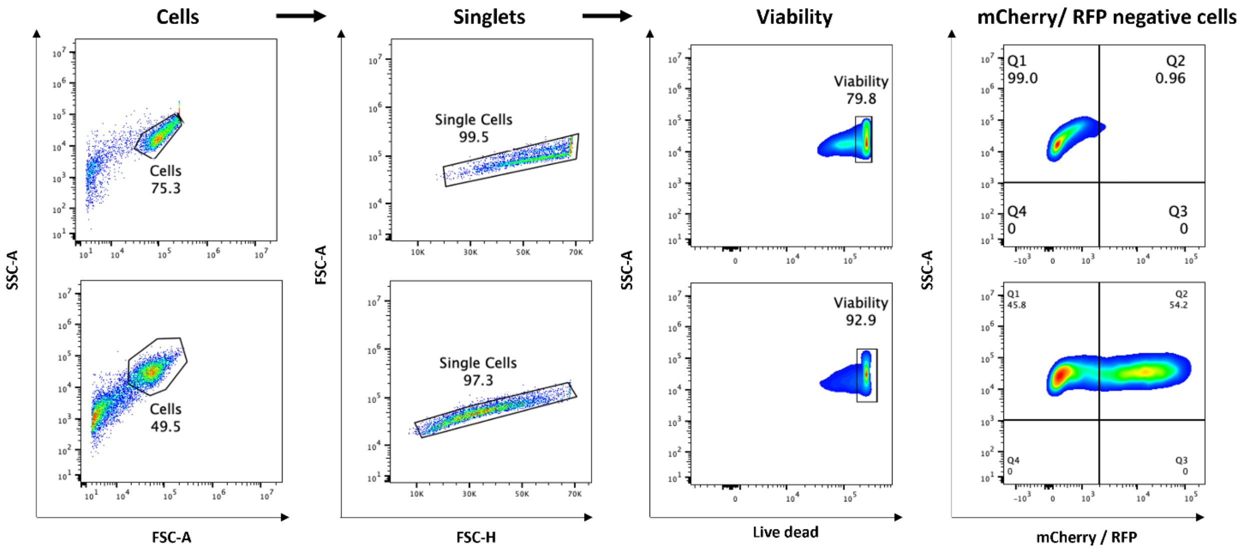


**Figure S8:** Gating strategy for the flow cytometry analysis. Single cell suspension was prepared from mock-infected (media without virus) and Pr8 and RSV infected cells. Cell suspensions were stained for viability and the viral infection was reported by mCherry. Proceeding from left to right, the initial dot plot depicted side scatter (SSC-A) on the Y-axis and forward scatter (FSC-A) on the X-axis. Applying gating to the cell population based on size-scatter enabled the visualization of individual cells in the dot plot featuring FSC-A (Y-axis) and FSC-H (X-axis). By gating on all singlet cell population, a dot plot representing SSC-A (Y axis) and viability marker (intensity mean, X axis) was generated, displaying cell viability by following the positive cells for the marker. Ultimately, by gating on all viable cell populations, we constructed a dot plot with SSC-A (Y-axis) and mCherry reporter (average intensity, X-axis). This allowed us to accurately determine the infection percentage within the given experimental condition.

**Pr8**

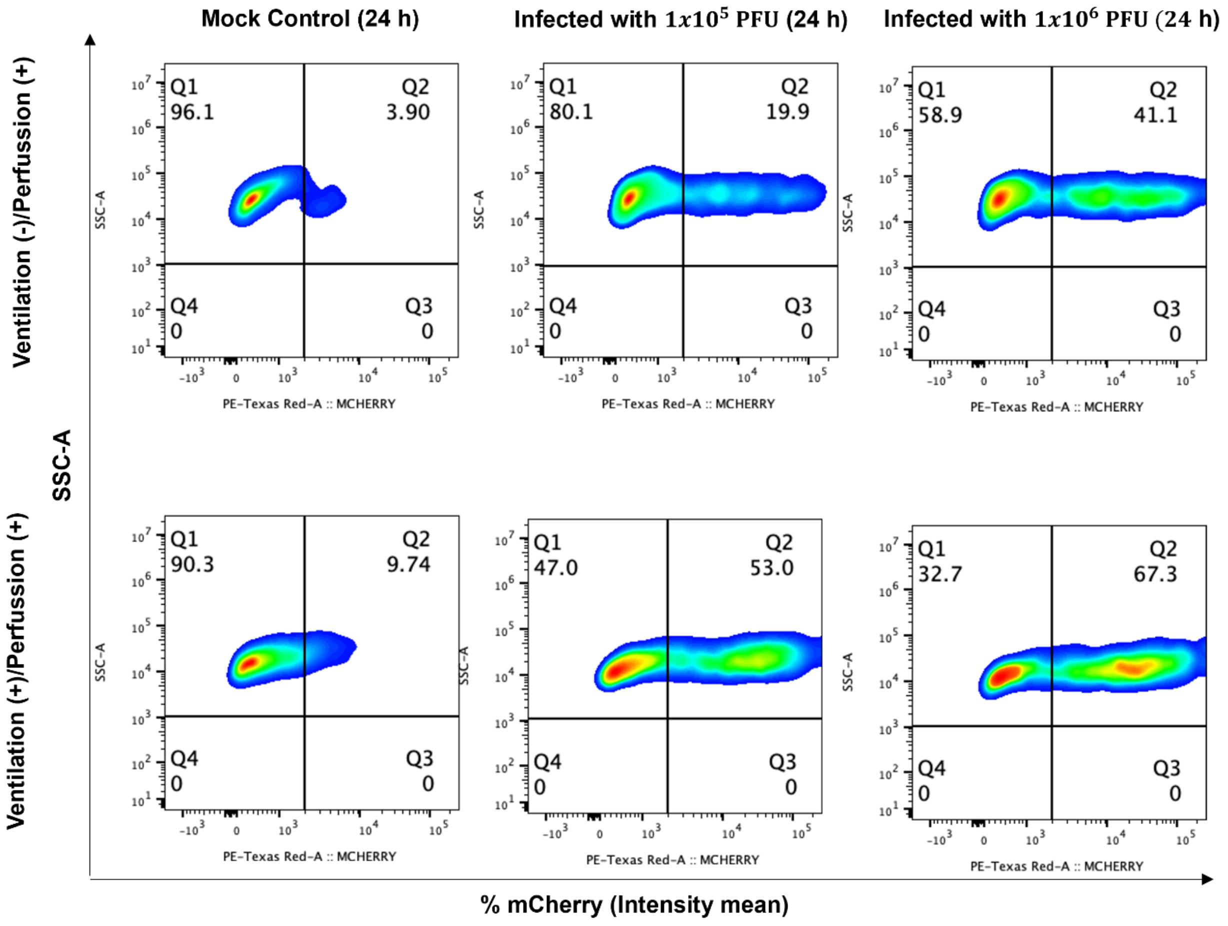


**Figure S9:** Representative histocytometry plots on 24 h Pr8 infected Ventilation (+)/Perfussion (+) and Ventilation (-)/Perfussion (+) groups. The mean staining intensity for cell populations was positive for Calcein (Y axis) and viral NP (X axis). The NP (for viral infection) channel was generated in Imaris 9.4 using the Channel Arithmetics Xtension prior to running surface creation to identify Calcein-NP cells in images. Statistics were exported for each surface and imported into FlowJo v10.3 for image analysis ($n$=3).

## Pr8

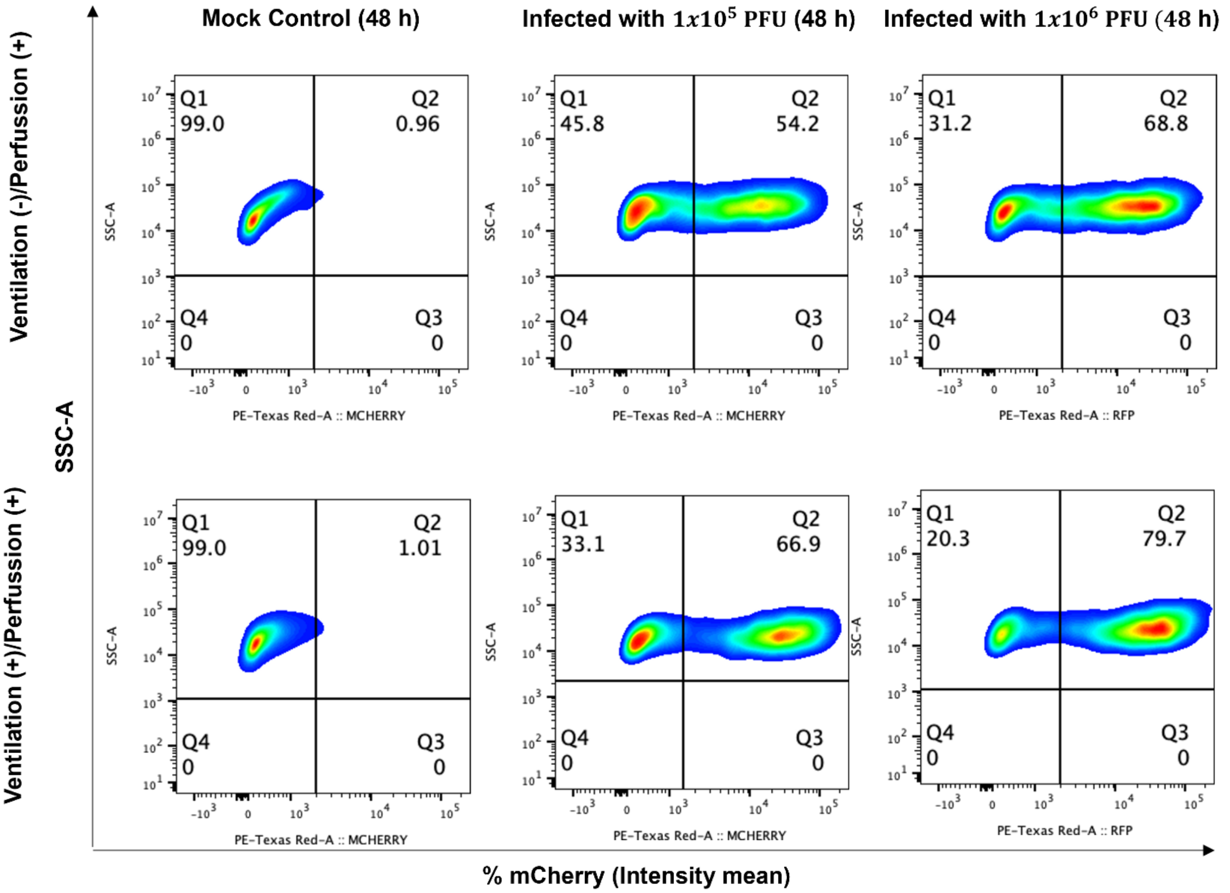


**Figure S10:** Representative histocytometry plots on 48 h Pr8 infected Ventilation (+)/Perfussion (+) and Ventilation (-)/Perfussion (+) groups. The mean staining intensity for cell populations was positive for Calcein (Y axis) and viral NP (X axis). The NP (for viral infection) channel was generated in Imaris 9.4 using the Channel Arithmetics Xtension prior to running surface creation to identify Calcein-NP cells in images. Statistics were exported for each surface and imported into FlowJo v10.3 for image analysis ($n$=3).

### RSV

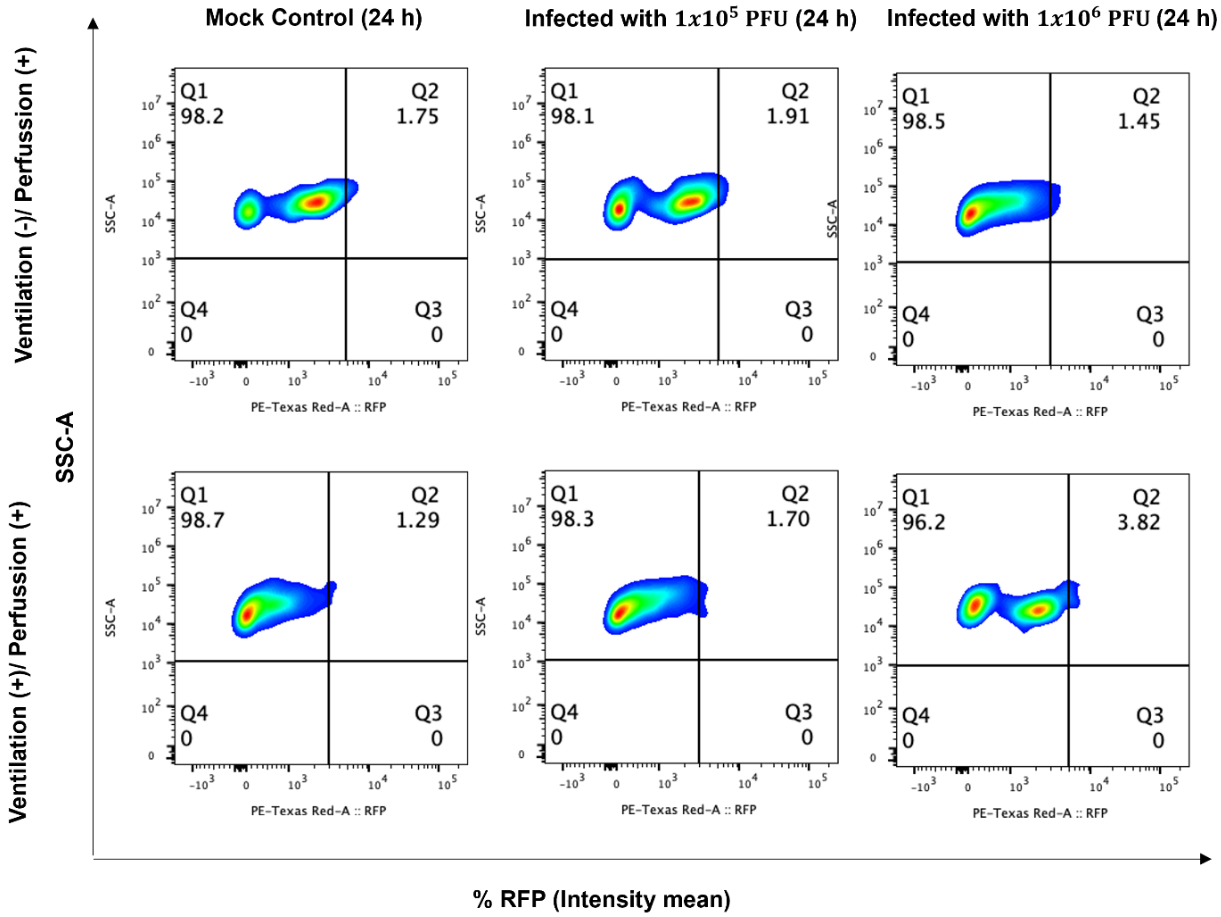


**Figure S11:** Representative histocytometry plots on 24 h RSV infected Ventilation (+)/Perfussion (+) and Ventilation (-)/Perfussion (+) groups. The mean staining intensity for cell populations was positive for Calcein (Y axis) and viral NP (X axis). The NP (for viral infection) channel was generated in Imaris 9.4 using the Channel Arithmetics Xtension prior to running surface creation to identify Calcein-NP cells in images. Statistics were exported for each surface and imported into FlowJo v10.3 for image analysis (*n*=3).

### RSV

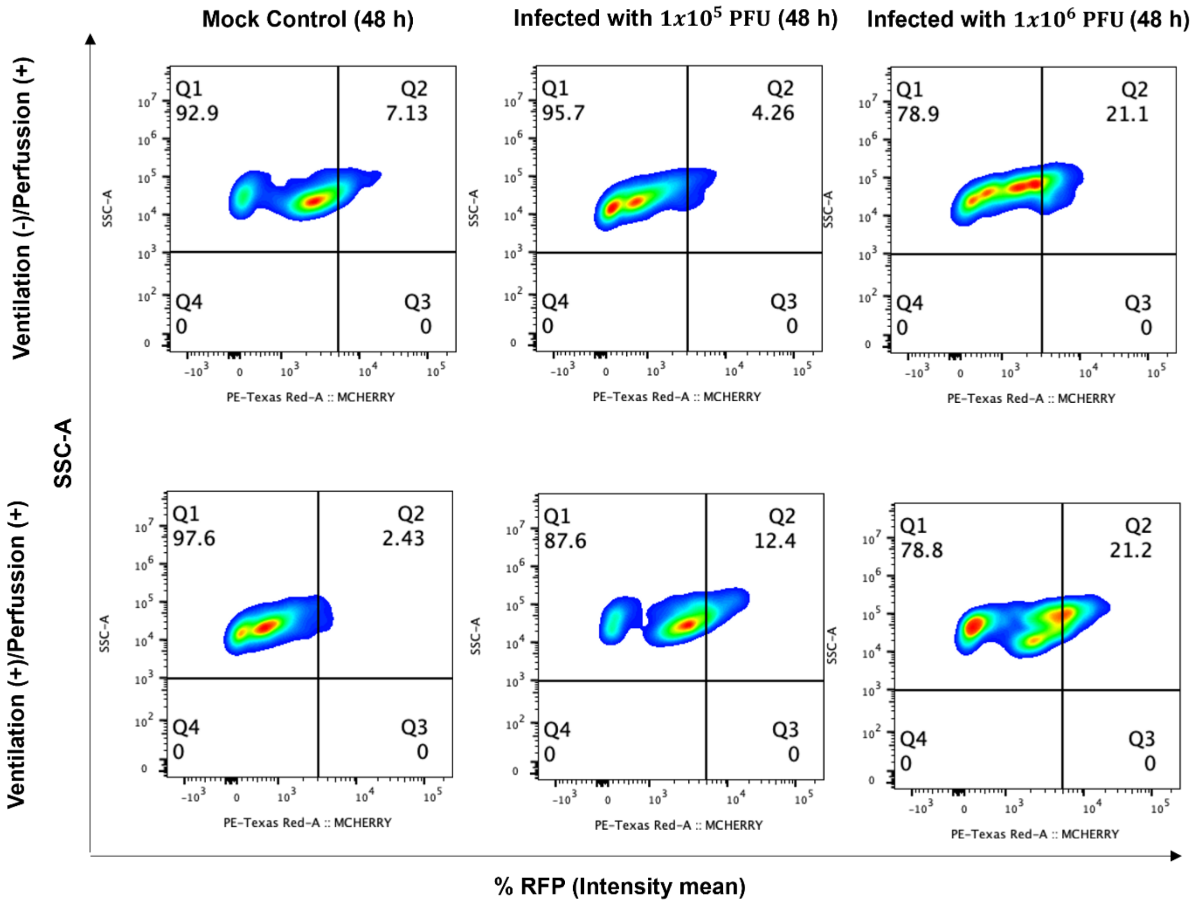


**Figure S12:** Representative histocytometry plots on 48 h RSV infected Ventilation (+)/ Perfussion (+) and Ventilation (-)/Perfussion (+) groups. The mean staining intensity for cell populations was positive for Calcein (Y axis) and viral NP (X axis). The NP (for viral infection) channel was generated in Imaris 9.4 using the Channel Arithmetics Xtension prior to running surface creation to identify Calcein-NP cells in images. Statistics were exported for each surface and imported into FlowJo v10.3 for image analysis ($n$=3).

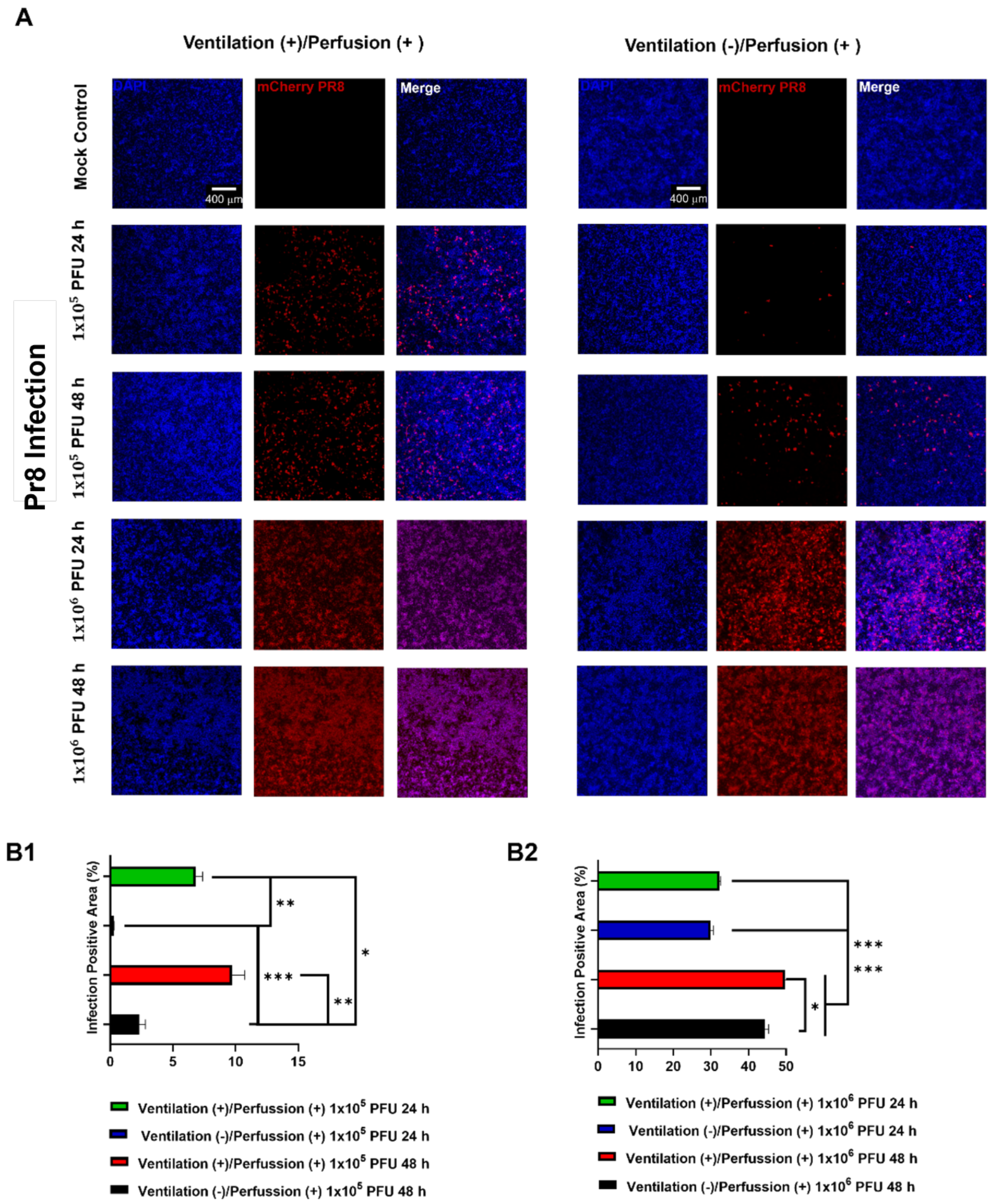


**Figure S13: (A)** Representative immunofluorescent images of Pr8 infection based on DAPI staining and NP positive signal. Cells were infected at different concentrations ($1x10^5$ or $1x10^6$ PFU) for 24 or 48 h. Quantification of Pr8 infection based on infection positive area. **(B1)** $1x10^5$ PFU for 24 or 48 h, **(B2)** $1x10^6$ PFU for 24 or 48 h ($n$=3; $p^{*}$<0.05, $p^{**}$<0.01 and $p^{***}$<0.001).

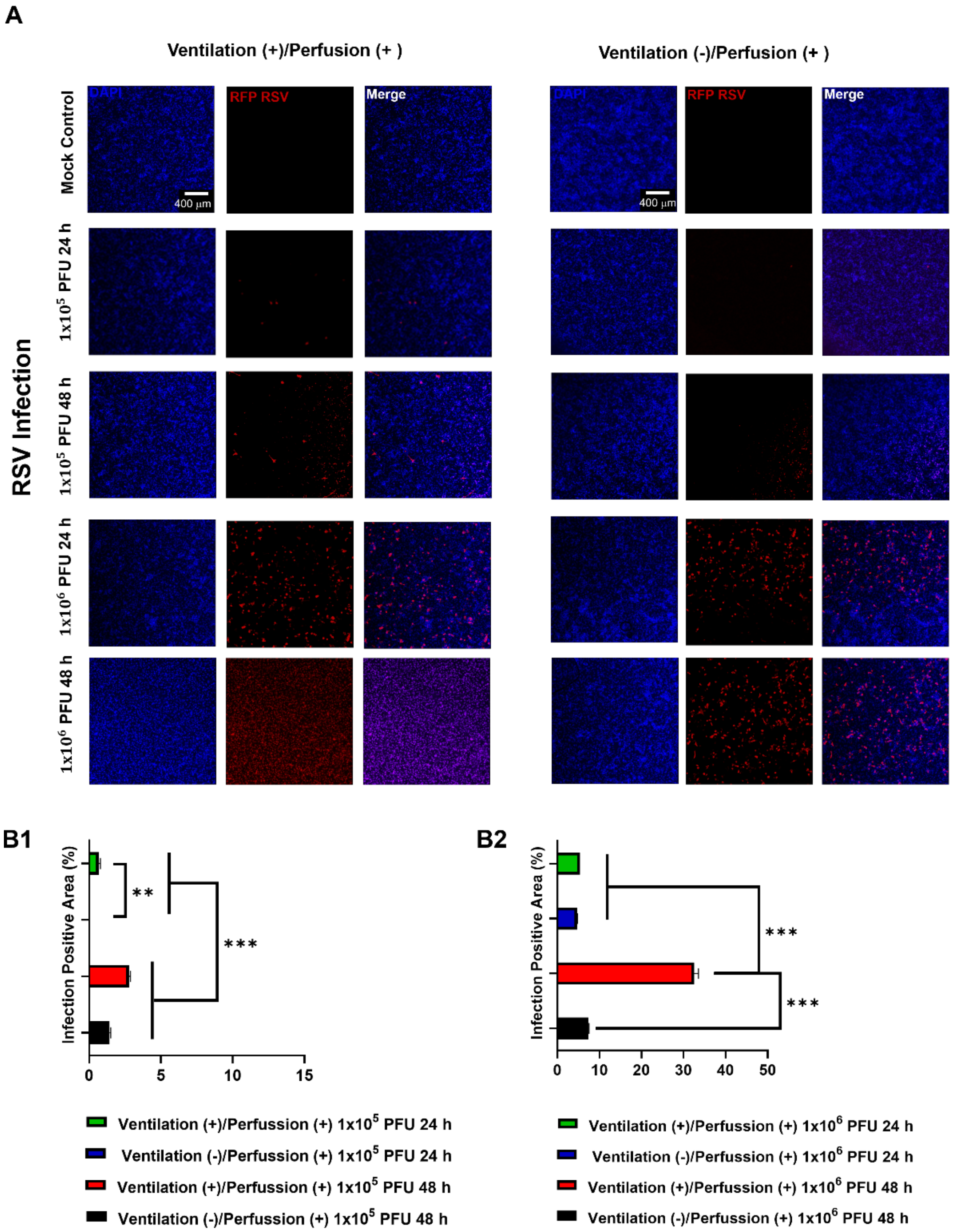


**Figure S14: (A)** Representative immunofluorescent images of RSV infection based on DAPI staining and NP positive signal. Cells were infected at different concentrations ($1x10^5$ or $1x10^6$ PFU) for 24 or 48 h. Quantification of RSV infection based on infection positive area. **(B1)** $1x10^5$ PFU for 24 or 48 h, **(B2)** $1x10^6$ PFU for 24 or 48 hf ($n$=3; $p^{**}$<0.01 and $p^{***}$<0.001).

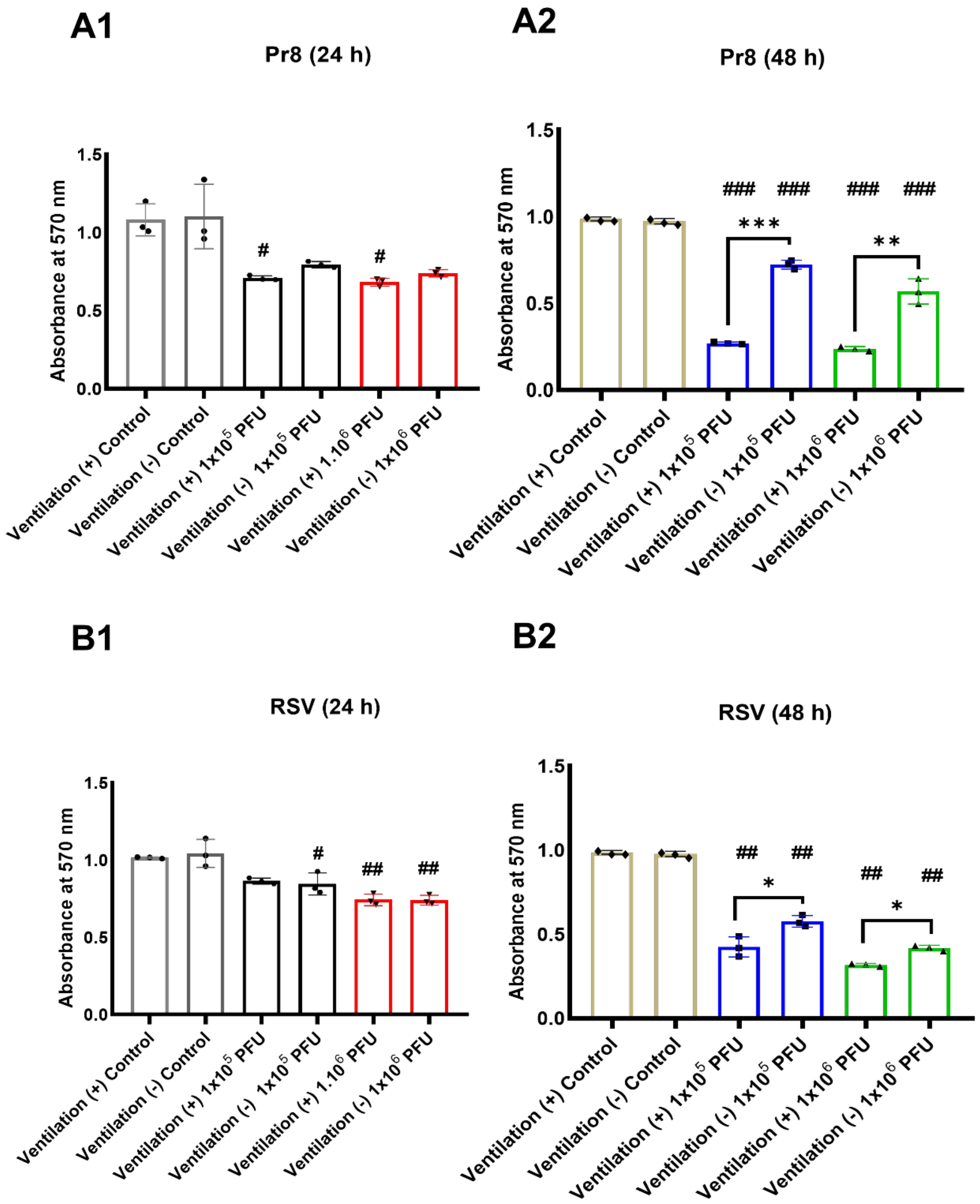


**Figure S15:** Comparative analysis of cell viability using the MTT assay. **(A1)** Pr8 Infection at 24 or **(A2)** 48 h, and RSV Infection at **(B1)** at 24 or **(B2)** 48 h of the viral infection ($n$=3; $^{*}$<0.05, $p^{**}$<0.01 and $p^{***}$<0.001; $p^{\#} < 0.05$, $p^{\#\#} < 0.01$ and $p^{\#\#\#} < 0.001$ indicate significance with respect to the control of the same group).

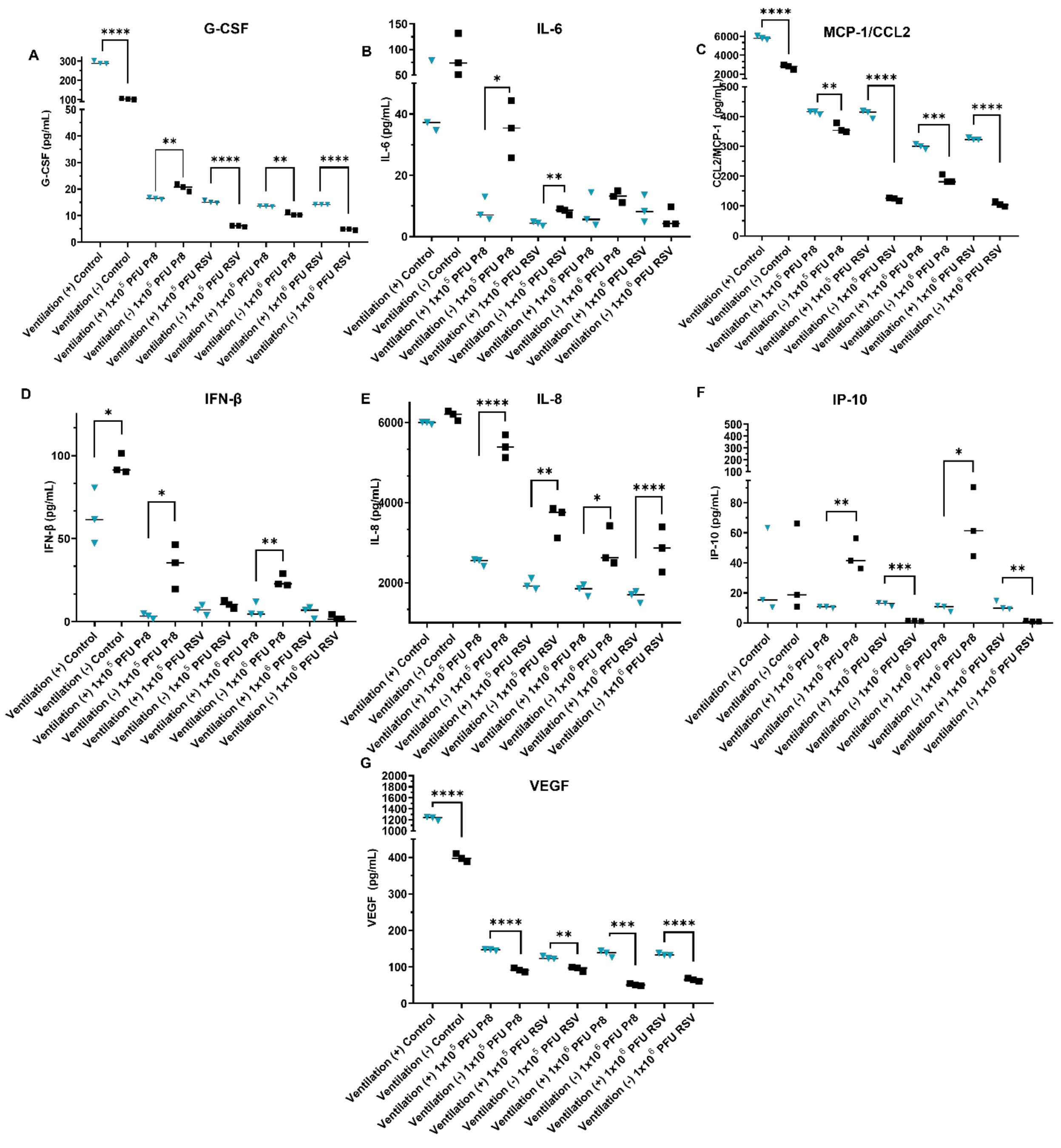


**Figure S16:** Lung device supernatant cytokines/chemokines measured using immunoassay across conditions. **(A)** G-CSF, **(B)** IL-6, **(C)** MCP-1/CCL2, **(D)** IFN-β**, (E)** IL-8**, (F)** IP-10, and **(G)** VEGF proteins measured in supernatants under all conditions at 24 h of Pr8 and RSV infections ($n$=3; $p^{*}$<0.05, $p^{**}$<0.01, $p^{***}$<0.001, $p^{****}$<0.0001).

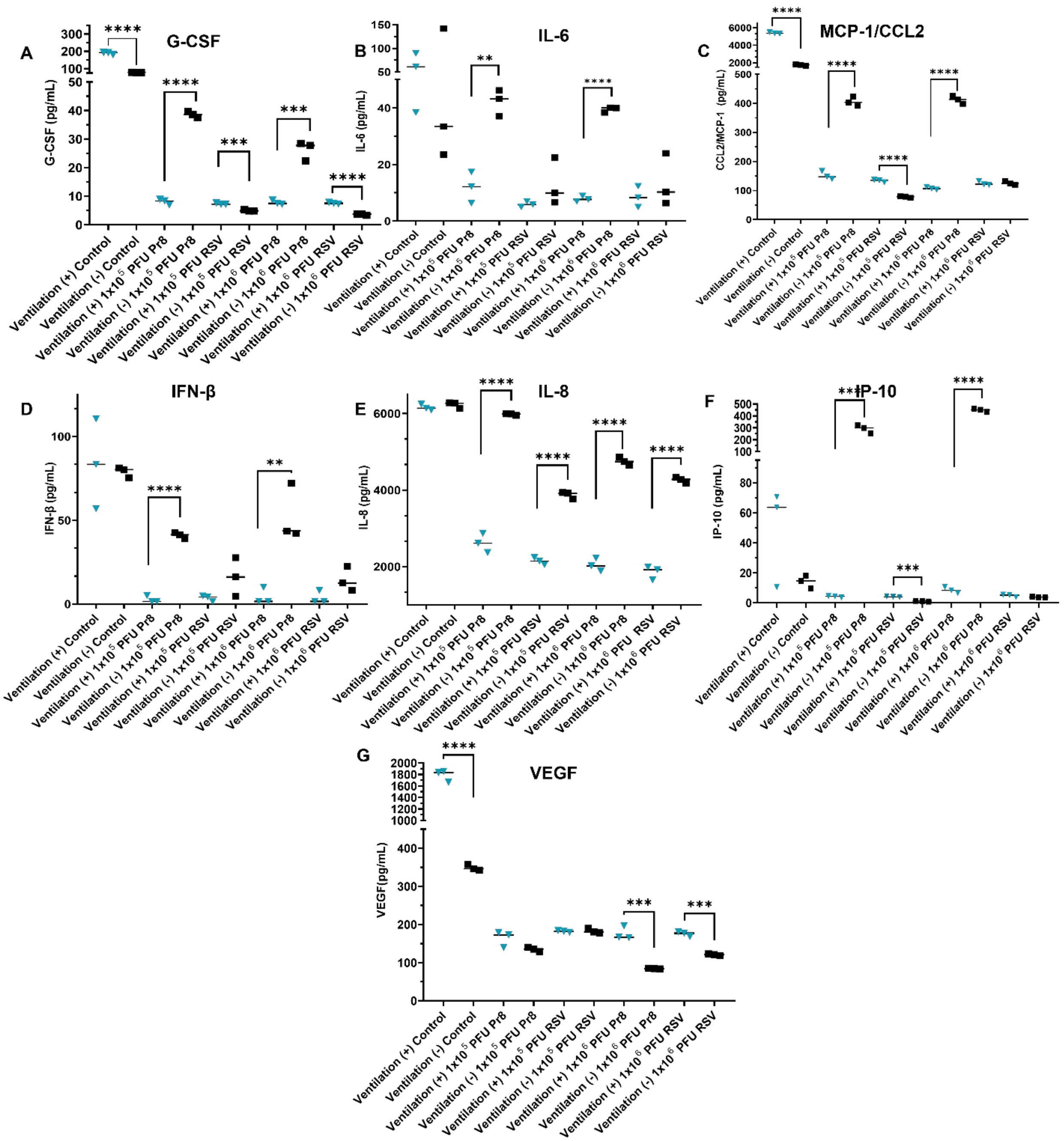

**Figure S17:** Lung device supernatant cytokines/chemokines measured using immunoassay across conditions. **(A)** G-CSF, **(B)** IL-6, **(C)** MCP-1/CCL2, **(D)** IFN-β**, (E)** IL-8**, (F)** IP-10, and **(G)** VEGF proteins measured in supernatants under all conditions at 48 h of Pr8 and RSV infections ($n$=3; $p^{*}<0.05$, $p^{**}<0.01$, $p^{***}<0.001$, $p^{****}<0.0001$).

**Video Captions**

**Video S1:** A representative 3D printed lung model platform under a continuous respiratory cycle and perfusion.
